# Distinct Temporal Responses to *Xanthomonas translucens* Strains Shape Bacterial Leaf Streak Development in Triticale

**DOI:** 10.64898/2026.09.07.749878

**Authors:** Fahad Hasan, Fazal Manan, Edward Cedrick Fernandez, Shuang Wu, Basanta Dhungana, Changhui Yan, Gongjun Shi, Zhaohui Liu, Zhikai Liang

## Abstract

Bacterial leaf streak caused by *Xanthomonas translucens* threatens cereal production, however, the temporal coordination of host transcriptional responses during resistant and susceptible interactions in polyploid crops remains partially understood. Here, we used time-resolved transcriptomics to characterize responses of synthetic hexaploid triticale to two *X. translucens* pv. *undulosa* strains that produce contrasting disease outcomes. The resistant interaction with non-virulent LB10 showed a strong early transcriptional response that subsequently declined, whereas responses to the virulent strain P3 progressively intensified as water-soaking symptoms developed. Analysis of syntenic A-, B-, and R-subgenome homoeologs revealed extensive regulatory asymmetry, with R-subgenome homoeologs disproportionately represented among transcriptionally suppressed genes. Despite their conserved coding sequences, homoeologs often showed divergent transcriptional responses during infection, whereas greater similarity in upstream regulatory regions was associated with more coordinated responsive trajectories. We next examined pathogen-mediated transcriptional regulation through transcription activator-like (TAL) effectors. Among eight TAL effector templates identified in LB10 and P3, TAL5-associated predicted targets showed the strongest preferential induction during P3 infection. Disruption of TAL5 in P3 predominantly reduced host gene expression, including genes involved in immune signaling, cell wall-associated defense and photosynthetic function, accompanied by reduced maximum photosystem II quantum efficiency at 72 h post-inoculation. Together, our results show that bacterial leaf streak outcomes are shaped by temporally distinct host responses and pathogen effector-associated transcriptional reprogramming, providing insight into the dynamic regulation underlying cereal-*Xanthomonas* interactions.

## Introduction

Bacterial leaf streak (BLS), caused by *Xanthomonas translucens*, is an important disease in cereal crops that can lead to substantial yield losses and poor grain quality under favorable environmental conditions (Sapkota et al., 2020; Osdaghi et al., 2023). The *Xanthomonas translucens* species complex comprises multiple pathovars with overlapping but distinct host ranges, including *pv. undulosa* (Xtu), which primarily infects common bread wheat, durum wheat and rye, while *pv. translucens* (Xtt), which is more commonly associated with barley (Bragard et al., 1997). Upon infection, *Xtu* colonizes the intercellular spaces of leaf tissues, leading to the development of characteristic disease symptoms (Roman-Reyna et al., 2024). In susceptible genotypes, early infection is often marked by water-soaked lesions that expand longitudinally along the leaf veins (Ledman et al., 2023) and progressively develop into chlorotic streaks and necrotic tissues (Murcia Bermudez et al., 2026). In contrast, resistant genotypes typically show limited or absent water-soaking and may develop chlorotic spots associated with restriction of bacterial proliferation (Sapkota et al., 2018; Acharya et al., 2024).

These outcomes, however, are not fixed states but the result of a dynamic process. Between initial infection and the appearance of visible symptoms, the host undergoes coordinated, time-dependent changes in pathogen recognition, defense signaling and metabolism (Castro-Moretti et al., 2020; Ngou et al., 2021; Wang et al., 2023). The timing and order in which these changes occur determine whether the infection ends in resistance or susceptibility. For example, in the rice-*Xanthomonas oryzae* pathosystem, resistant plants activate defense pathways sequentially, with early phytoalexin and jasmonic acid/salicylic acid signaling followed by cell wall reinforcement, emphasizing the importance of temporal coordination in disease resistance (Zhang et al., 2025). Consistent with this, early transcriptional divergence between compatible and incompatible interactions has also been observed in tomato challenged with pathogens, where divergence in gene expression was detectable well before differences in visible symptom progression became apparent (Rezzonico et al., 2017), underscoring that infection outcome was often set well before phenotypic symptoms emerge. In cereals infected with *X. translucens*, transcriptomic and proteomic profiling of wheat has identified defense-associated responses, including jasmonic acid and hydrogen peroxide signaling (Garcia-Seco et al., 2017). However, whether and how these host responses are temporally coordinated over the course of *X. translucens* infection remains uncharacterized.

Part of this temporal reprogramming is driven directly by the pathogen. A defining feature of *Xanthomonas* pathogenesis is the deployment of transcription activator-like (TAL) effectors, which recognize specific effector binding elements (EBEs) in host promoters through arrays of repeat variable diresidues (RVDs) to activate target gene expression (Ji et al., 2016; Teper et al., 2023). Because TAL effectors act by directly inducing transcription of their targets, their activity imposes a specific temporal signature on host gene expression – activating susceptibility genes to promote disease, or triggering executor resistance genes to activate host immunity (White et al., 2009; Shantharaj et al., 2017; Xu et al., 2017; Teper et al., 2023). In the well-characterized rice-*Xanthomonas oryzae* pathosystem, TAL effectors target susceptibility genes such as members of the *SWEET* family (Oliva et al., 2019; Xu et al., 2024). Similarly, TAL effectors have been implicated in *X. translucens* virulence in cereals (Gutierrez-Castillo et al., 2024), including a TAL effector from *X. translucens* pv. *undulosa* that promotes disease progression by inducing *TaNCED-5BS* expression and altering ABA signaling in wheat (Peng et al., 2019). Yet computational tools for TAL effector target prediction, such as AnnoTALE (Grau et al., 2016) and PrediTALE (Erkes et al., 2019) which were developed and benchmarked primarily on rice-*Xanthomonas oryzae* data. Whether these prediction frameworks could be translated to other cereal-*Xanthomonas* systems, including *X. translucens*, remains not well understood.

This question is further complicated in polyploid crops, where multiple homoeologous gene copies distributed across subgenomes exhibit distinct transcriptional behaviors (He et al., 2022). Previous studies have shown that subgenomes can contribute unequally to gene expression (Bird et al., 2018), with homoeologs exhibiting biased or dynamically partitioned expression patterns under stress conditions (Dong and Adams, 2011; Powell et al., 2017; Lee and Adams, 2020; de Jong and Adams, 2023; Peng et al., 2024). These patterns indicate that stress-responsive regulation is not necessarily coordinated among homoeologs and may instead reflect differences in landscapes of their underlying regulatory sequences (Ramírez-González et al., 2018). Such regulatory divergence creates an additional layer of complexity during pathogen infection, particularly when pathogen-derived signals act through specific promoter sequences. In this context, sequence variation among homoeologous promoters can result in differential responsiveness even among otherwise closely related gene copies. However, how variations in regulatory sequences contribute to subgenome-specific transcriptional responses during bacterial infection still need to be explored, particularly in polyploid cereal-*Xanthomonas* interactions.

Triticale (x *Triticosecale spp*.), a synthetic allohexaploid formed through hybridization between durum wheat (*Triticum turgidum subsp. durum*; A and B genomes) and rye (*Secale cereale*; R genome), provides a system to investigate BLS infection in a polyploid context. Due to its hybrid origin, triticale retains substantial subgenome divergence, particularly between the wheat-derived A/B genomes and the rye-derived R genome (Ma and Gustafson, 2008). This divergence creates a natural context in which identical pathogen effectors encounter distinct promoter landscapes across homoeologs, enabling assessment of how *cis*-regulatory variation influences subgenome-specific transcriptional responses. Here, we performed time-resolved transcriptomic profiling of triticale infected with two *Xtu* strains, LB10 and P3, which produce contrasting disease outcomes (Sapkota et al., 2018; Wen et al., 2018), to dissect how host and pathogen-driven transcriptional programs unfold over the course of infection. By tracking gene expression across multiple time points and comparing strains with divergent virulence, we characterize the temporal sequence of defense activation, subgenome-specific expression dynamics and TAL-target induction that together distinguish resistance from susceptibility outcomes in a polyploid cereal host.

## Results

### Time-resolved response divergence in triticale during infection by LB10 and P3

Two *Xtu* strains, LB10 (non-virulent) and P3 (virulent), induced different reactions in the hexaploid triticale cultivar ‘Siskiyou’ over time (Figure 1A). There was no visible reaction at the infiltrated spots 24 hr post-infiltration (hpi) for both strains. However, the virulent strain P3 produced a progressively intensifying water-soaking symptom from 48 hpi. In contrast, LB10 induced no visible water-soaking reaction till 96 hpi when a clear chlorosis developed within the infiltrated area, suggesting an incompatible and resistant interaction (Figure 1A). Genetic analysis and mapping have demonstrated that Syskiyou carries a dominant resistance gene (Xct1) on chromosome 5R against LB10 (Wen et al. 2017). To capture distinct temporal transcriptional responses, leaf tissues were collected in three replicates at four time points (24, 48, 72, and 96 hpi) after the inoculation with two strains as well as corresponding samples from mock inoculation with buffer alone, which yielded a total of 36 RNA-seq libraries. RNA-seq reads were aligned to a synthetic triticale reference transcriptome assembled from the durum wheat and rye cDNA sequences and 70,131 genes with mean CPM (counts per million) >1 across all samples were identified. Principal component analysis separated control, LB10, and P3 samples, with 24 hpi samples forming a distinct transcriptional state from later time points (Figure 1B), suggesting distinct early regulatory reprogramming.

**Figure 1.**
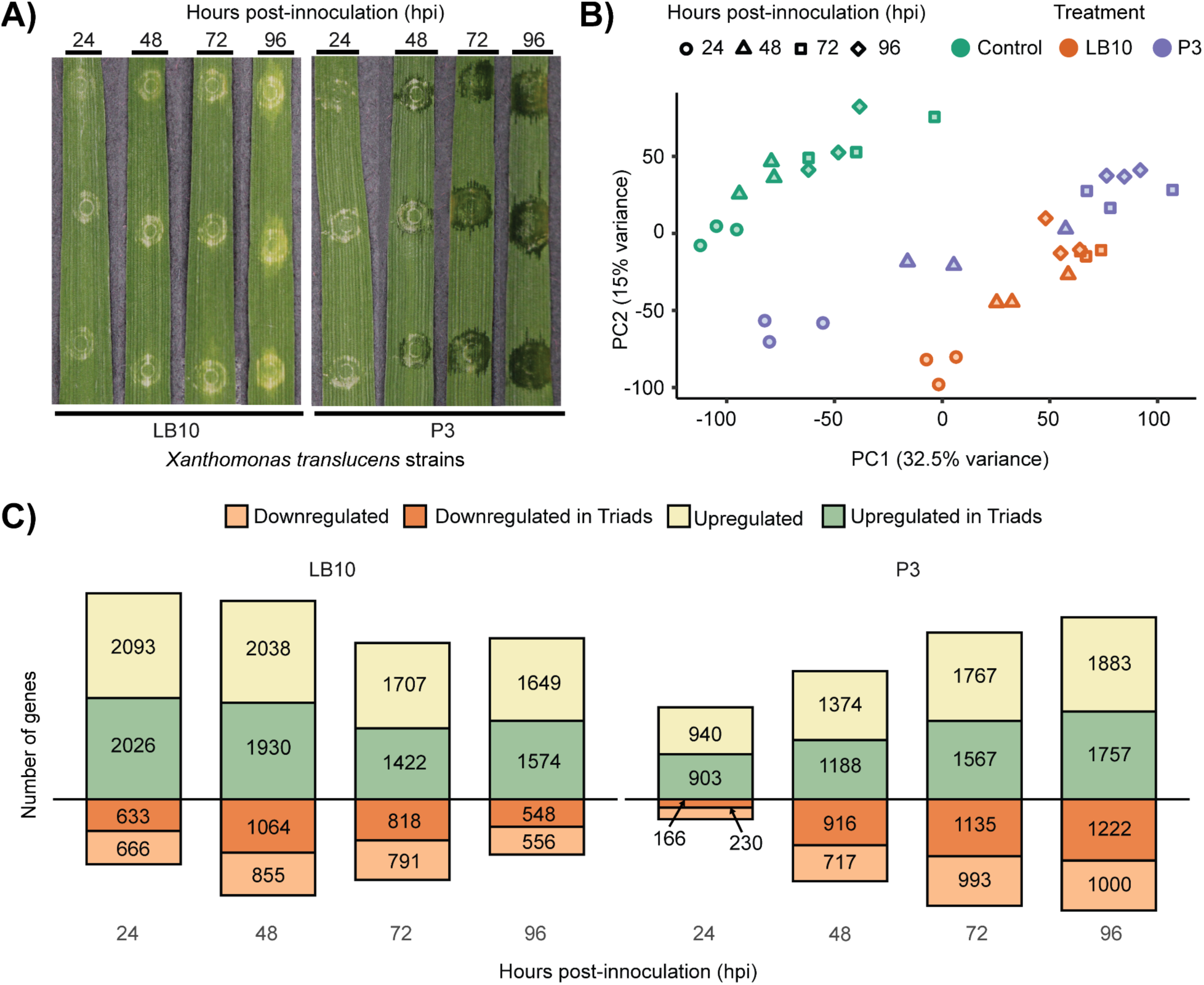
Temporal phenotypic and transcriptional responses of triticale to two *Xanthomonas translucens* pv. *undulosa* strains. (A) Leaf symptoms following inoculation with LB10 (resistant interaction) and P3 (susceptible interaction) at 24, 48, 72, and 96 hours post-inoculation (hpi); (B) Principal component analysis of normalized RNA-seq expression showing separation of samples by treatment and time point; (C) Numbers of DEGs under LB10 and P3 infection across time points, categorized by direction of regulation and whether they were part of syntenic homoeolog triads.

To determine whether LB10 and P3 induced distinct temporal transcriptional responses in triticale, we compared gene expression between inoculated and control plants at each time point. LB10 inoculation resulted in 5,418, 5,887, 4,738, and 4,327 differentially expressed genes (DEGs) at 24, 48, 72, and 96 hpi, respectively. In contrast, the number of DEGs under P3 inoculation progressively increased over time, from 2,239 at 24 hpi to 4,195, 5,462, and 5,862 at 48, 72, and 96 hpi, respectively (Figure 1C). These contrasting patterns indicated that LB10 elicited a relatively strong early transcriptional response, whereas the response to P3 intensified during disease progression. Because the A, B, and R subgenomes collectively constitute the synthetic hexaploid genome of triticale, we identified syntenic homoeologous groups across the three subgenomes. A strict homoeolog triad was defined as a group containing one syntenic gene from each subgenome. Of 13,112 generated syntenic homoeolog triads, 12,857 showed a strict 1:1:1 correspondence among A-, B-, and R-subgenome genes. The remaining 255 triads displayed one-to-many or many-to-many relationships in at least one subgenome, representing over 1.94% of all 13,112 triads and suggesting potential subgenome-specific divergence within triticale. Across strain-control DEGs per time point, approximately 50% were associated with syntenic homoeolog triads (Figure 1C).

Given the distinct phenotypic outcomes of LB10 (non-virulent) and P3 (virulent) inoculations, we compared enriched GO terms between these two infections. In P3, photosynthesis-related terms, including “photosynthesis, light harvesting” (GO:0009765) and “photosystem II” (GO:0009523), were consistently enriched relative to LB10 across the entire time course, with the divergence becoming most pronounced at 72-96 hpi, indicating a more sustained disruption of photosynthetic function in the susceptible interaction. Stomatal complex morphogenesis (GO:0010103) was additionally enriched in P3 at 72 and 96 hpi. In LB10, uniquely enriched terms instead pointed toward defense-associated functions in a temporally ordered manner, such as “endochitinase activity” (GO:0008843, PR-3 family) at 48 hpi, “protein phosphorylation” (GO:0006468) and “ADP binding” (GO:0043531) at 72 hpi – categories that encompass receptor kinases and the nucleotide-binding domains of NLR-type resistance proteins respectively (Tameling et al., 2002), and “glucan endo-1,3-β-D-glucosidase activity” (GO:0042973, PR-2 family) at 96 hpi (van Loon et al., 2006), indicating that defense gene activity was maintained in the resistant interaction throughout infection (Supplementary Table 1).

### Subgenome-specific gene contributions to divergent responses between LB10 and P3

As a synthetic hexaploid, triticale might coordinate BLS responses through syntenic homoeologs across the A, B, and R subgenomes. Based on the relative expression of three homoeologs within each triad, syntenic triads were classified into seven homoeolog expression-bias categories for each time point (see Methods; Figure 2A). Triad categories were more consistent over time within the same condition than between conditions at the same time point (Figure 2B). This trend was similarly observed for triad expression dynamics across all time points and conditions (Supplementary Figure 1). Because the R subgenome originated from rye and was combined with the A and B subgenomes of durum wheat during triticale synthesis, we asked whether homoeolog expression was balanced among the three subgenomes or showed subgenome-specific bias. Syntenic triads were assigned more frequently as recessive than dominant across all samples. Dominant genes were approximately evenly distributed among the A, B, and R subgenomes (A:B:R ≈ 1:1:1), although slightly fewer were assigned to the R subgenome. Intriguingly, recessive genes showed a pronounced bias toward the R subgenome (A:B:R ≈ 1:1:3), a pattern that was consistent across time points and treatments (Figure 2B; Supplementary Figure 1). However, this bias was not observed in bread wheat, where dominant and recessive homoeologs were distributed approximately equally across the A, B, and D subgenomes (Ramírez-González et al., 2018).

**Figure 2.**
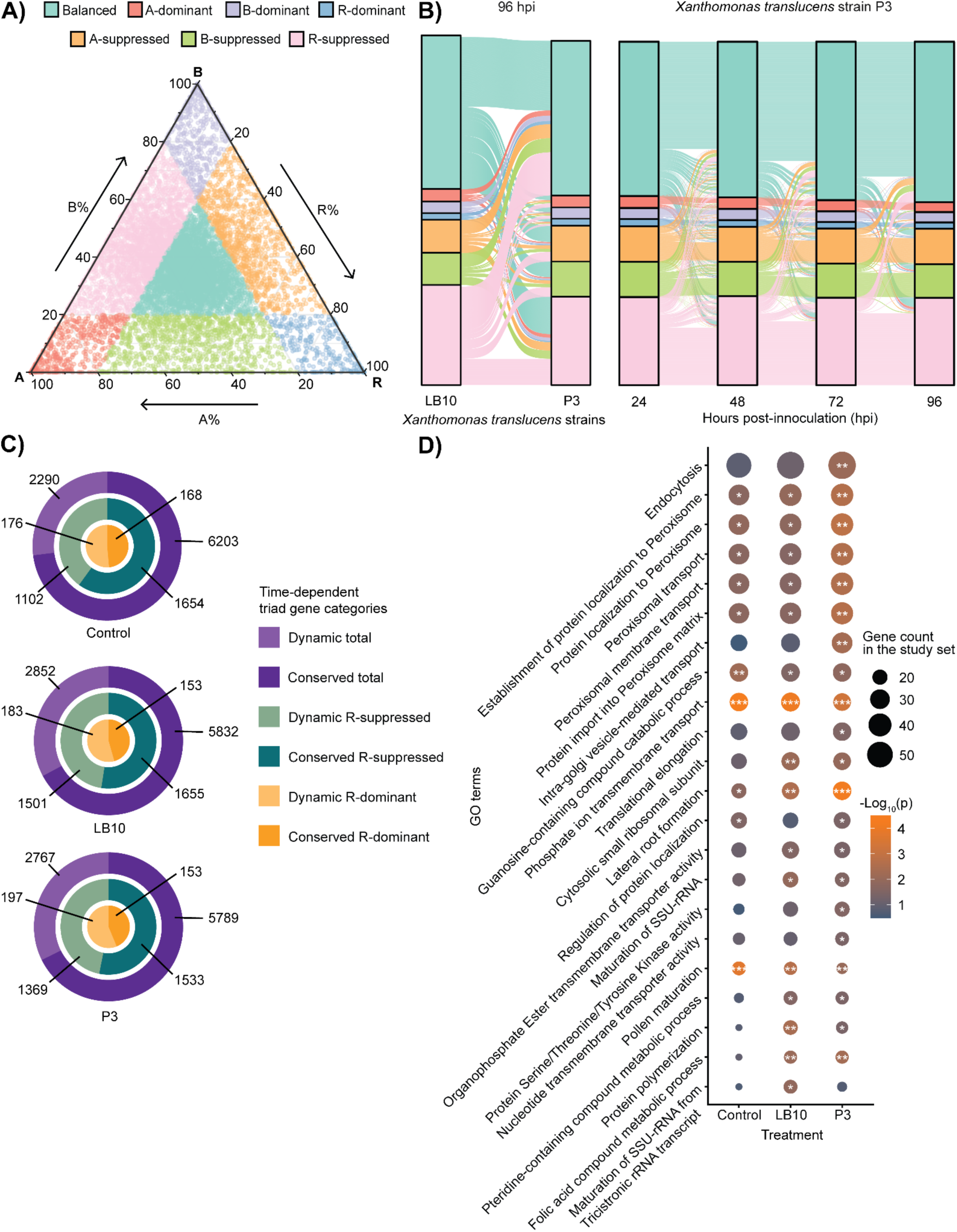
Temporal dynamics of syntenic homoeolog triad expression in triticale during BLS development. (A) Classification of syntenic homoeolog triads based on the relative expression contributions of the A-, B- and R-subgenome homoeologs, illustrated using P3 data at 48 hpi. Each point represents one triad, and its position reflects the relative contribution of each homoeolog to total expression in a triad; (B) Changes in triad expression categories between LB10 and P3 at 96 hpi (left) and across 24, 48, 72, and 96 hpi under P3 infection (right). Corresponding temporal patterns for other conditions were shown in Supplementary Figure 1; (C) Temporal conservation of triad expression categories under control, LB10, and P3 conditions. Outer rings represent all triads, middle rings represent R-suppressed triads, and inner rings represent R-dominant triads; (D) GO enrichment of R-subgenome genes from R-suppressed triads under control, LB10, and P3 conditions. Only GO terms significantly enriched in at least one condition are shown. GO terms with study counts >15 were retained, and the top 22 terms were ranked by their average enrichment ratio across the three conditions. Asterisks indicate adjusted p-values: * < 0.05, ** < 0.01, and *** < 0.001.

To examine the temporal stability of triads within each condition, we classified them as either conserved if their expression category remained consistent across all time points, or dynamic if their category changed over time. Overall, ∼72% triads exhibited conserved patterns rather than dynamic changes. Within this framework, R-suppressed triads exhibited a more balanced distribution between conserved and dynamic patterns, whereas R-dominant triads showed a slight enrichment in dynamic patterns relative to conserved ones (Figure 2C). Among 12,587 triads, 137 shared identical R homoelogs but were paired with different A and/or B homoelogs. However, only 19-24 of these showed inconsistent classifications, indicating that such cases had minimal impact on the overall proportions of dynamic and conserved triads.

Given the distinct evolutionary origin of the R subgenome relative to the wheat-derived A and B subgenomes, we next asked whether R-dominant or R-suppressed homoelogs were associated with specific biological functions during infection. Initial GO enrichment analyses were performed separately at each time point but only a few significant terms were identified, likely because relatively few genes were classified as R-dominant or R-suppressed at individual time points. Instead, to capture functions associated with R-subgenome expression bias across all stages, we focused on union sets of R-subgenome genes by pooling those assigned to R-suppressed or R-dominant triads across four time points. Using these union sets, no GO terms were significantly enriched among R-dominant genes, likely reflecting the substantially smaller number of genes in this category (Control: 356; LB10: 453; P3: 483) compared with R-suppressed genes (Control: 2,890; LB10: 3,467; P3: 3,260). In contrast, 75 GO terms were significantly enriched among R-suppressed genes in at least one treatment (Supplementary Table 2). Many enriched terms were related to nutrient and metabolite transport and were shared across control, LB10, and P3 treatments. Among these, “phosphate ion transmembrane transport” (GO:0035435) was the most significantly enriched term in all three treatments (Figure 2D), suggesting that suppression of R-subgenome homoeologs was broadly associated with transport and homeostatic functions. A smaller subset of enriched terms was treatment specific and potentially related to pathogen-associated processes. For example, P3-specific enrichment included “endocytosis” (GO:0006897), whereas LB10-specific enrichment included protein “K63-linked ubiquitination” (GO:0070534). Together, these results indicated that R-suppressed homoeologs were predominantly associated with conserved transport and regulatory functions, while a subset showed treatment-specific functional enrichment during pathogen infection.

### Temporal clustering reveals homoeolog regulatory divergence during BLS infection

To characterize strain-specific transcriptional dynamics during BLS infection, we performed k-means clustering using standardized log_2_fold-change values from LB10 and P3 across four time points (24, 48, 72, and 96 hpi). Joint clustering of genes in response to both strains allowed genes to be grouped according to their temporal responses while directly capturing response differences between LB10 and P3. We identified 16 temporal response clusters (see Methods; Supplementary Figure 2), revealing distinct transcriptional trajectories across LB10 and P3 (Figure 3A). To facilitate functional interpretation and comparison among clusters, semantically related enriched GO terms were consolidated into representative functional categories (Supplementary Table 3). Across the 16 temporal clusters, enriched GO terms fell into recurring functional themes, with two clusters showing particularly clear strain-divergent temporal patterns. Cluster 10 was distinctly enriched for antimicrobial and cell-death-associated representative functions (killing of cells of another organism; stress-induced premature senescence), with a temporal profile that diverged sharply by strain: LB10 showed early, elevated expression that declined steadily over the time course, whereas P3 rose only gradually to a later, lower peak – consistent with a more rapid and transient antimicrobial/cell-death response in the resistant genotype. Cluster 2, enriched for photosystem-associated terms (photosynthesis/PSII light harvesting), showed a strain-divergent temporal profile in which P3 declined sharply from an early peak while LB10 remained comparatively stable throughout the time course — consistent with progressively developed water-soaking symptoms in P3. Distinct transcription factor binding motifs were enriched among individual temporal clusters (Supplementary Table 4). The most robust signal was a GCC-box-like element (CCGCCGCCGCCR), which was enriched in cluster 1, 5, 9 and 11, consistent with the established role of GCC-box/ERF-mediated signaling in pathogenesis-related gene activation (Ohme-Takagi and Shinshi, 1995). Cluster 7, marked by a transient LB10 peak and a late decline in both strains toward the final time point, was enriched for a MYB/MYB-related-associated element (TGGATAAGG), a family with documented roles in pathogen defense transcriptional regulation (Raffaele and Rivas, 2013).

**Figure 3.**
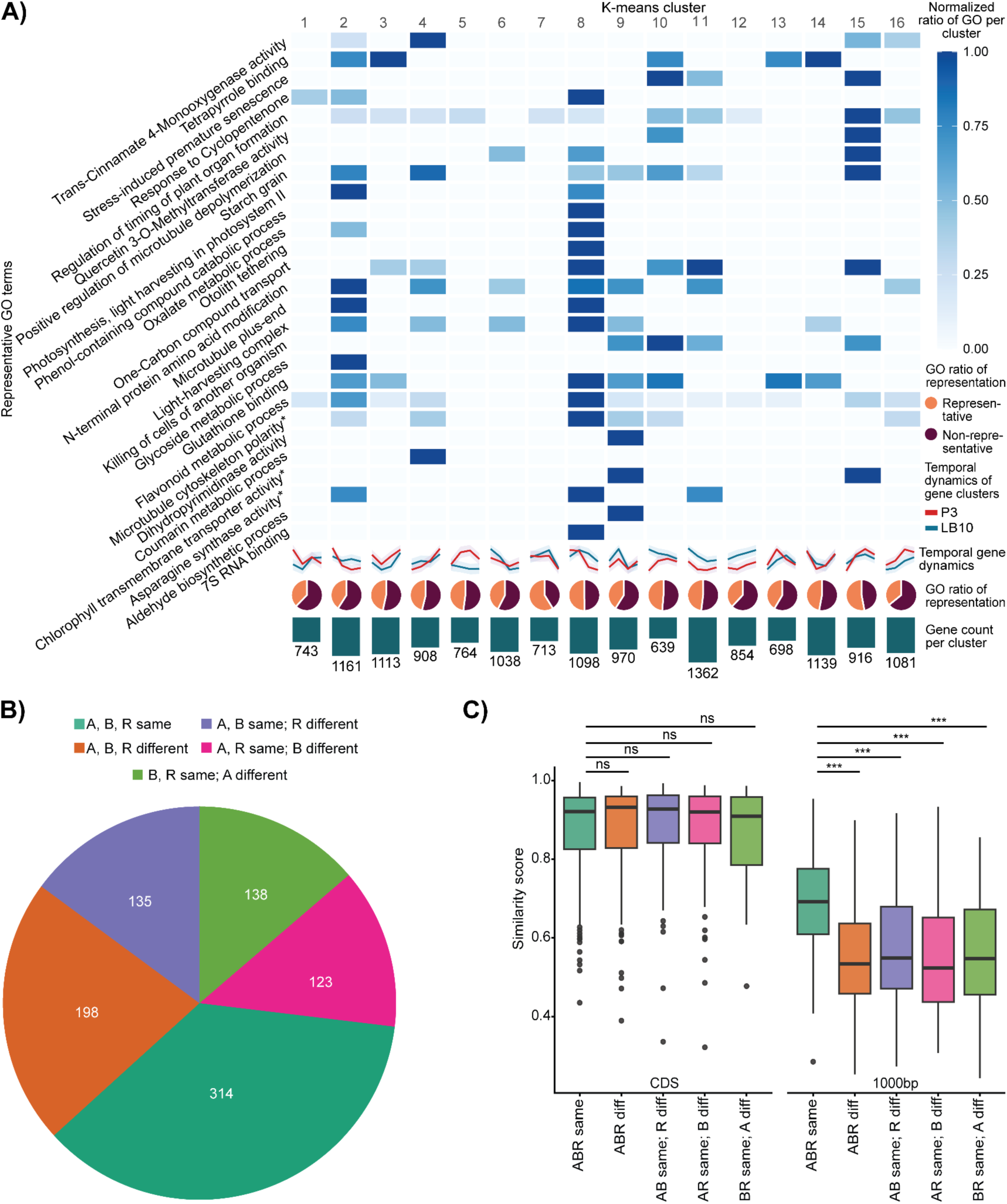
Functional clustering and BLS response divergence among homoeologs in triticale. (A) Expressed genes were grouped into 16 clusters based on their temporal response patterns using K-means clustering. GO enrichment analysis was performed for each cluster, and significantly enriched GO terms were consolidated into representative functional categories. Only categories comprising more than two individual GO terms were included in the heatmap. For each category, the proportion of associated GO terms was normalized across the 16 clusters. Asterisks (*) indicate abbreviated category names used for presentation. The complete GO enrichment results are provided in Supplementary Table 3; (B) Distribution of syntenic homoeolog triads according to how their homoeologs were assigned to the same cluster; (C) Pairwise sequence similarity of coding sequences and 1-kb upstream regions among homoeologs within triads, grouped by clustering pattern. Statistical significance is indicated by ns, not significant, and ***, p < 0.001.

Beyond these host *cis*-regulatory elements recognized by endogenous transcription factors, *Xanthomonas* could also directly hijack host gene expression through TAL (Transcription Activator-Like) effector that binds promoter sequences to activate susceptibility genes. TAL effectors were known to activate host susceptibility genes through SWEET family sugar transporters in rice (Streubel et al., 2013). Among 102 candidate SWEET genes (Supplementary Table 5), we identified 9 and 16 DEGs in P3 and LB10 across four time points, respectively. However, none of the SWEET genes exhibiting differential expression were predicted targets of any TAL effector template. As homoelogs within a same triad often retained highly similar sequences, we asked whether they also exhibited coordinated transcriptional responses during infection or diverged in their temporal regulation. Among the 15,197 genes represented across 16 clusters, 2,724 (17.92%) of them were assigned to three homoeologs in 908 syntenic triads. Of these, 314 triads (35%) had all three homoeologs assigned to the same cluster, 396 (43%) had two homoeologs co-clustering while the third diverged, and 198 (22%) showed no two homoeologs in a common cluster (Figure 3B). To test whether sequence conservation explained divergent cluster assignments, we compared CDS and 1kb upstream promoter similarity among A, B, and R syntenic homoelogs (see Methods). As expected, CDS similarity was comparable between triads with co-clustered and divergently clustered homoelogs (adjusted p-value >= 0.64), indicating that coding sequence conservation did not explain transcriptional coordination. In contrast, promoter similarity was significantly higher among triads with fully or partially co-clustered homoelogs than among those with divergent cluster assignments (adjusted p-value <= 3.91 x 10^-11^). These results suggested that sequence variation in upstream regulatory regions might contribute to divergent BLS-responsive expression patterns among homoelogs (Figure 3C).

### Effects of predicted TAL effector binding sites on host transcriptional responses to BLS

*Xanthomonas translucens* deploys TAL effectors that function as pathogen-delivered transcription factors, providing an opportunity to investigate host genes directly targeted by effector-mediated transcriptional activation. To identify potential TAL effector targets of LB10 and P3, we therefore analyzed their genome sequences using AnnoTALE (Grau et al., 2016) to characterize TAL effector RVD arrays and predict the corresponding host effector binding elements (EBEs). Among the 8 TALE RVDs identified per strain, only two RVD pairs exhibited differences between LB10 and P3. One pair (TAL template 1; TAL1) exhibited a three-RVD truncation in P3 compared with its counterpart in LB10 (Figure 4A). Another pair (TAL template 5; TAL5) only differed by a single RVD mismatch, where the 11th RVD was NI in LB10 and NN in P3, while all remaining 14 RVDs were identical (Figure 4A). NI preferentially recognizes adenine, whereas NN recognizes both adenine and guanine, suggesting a difference in nucleotide-recognition specificity between the LB10 and P3 TAL template 5.

**Figure 4.**
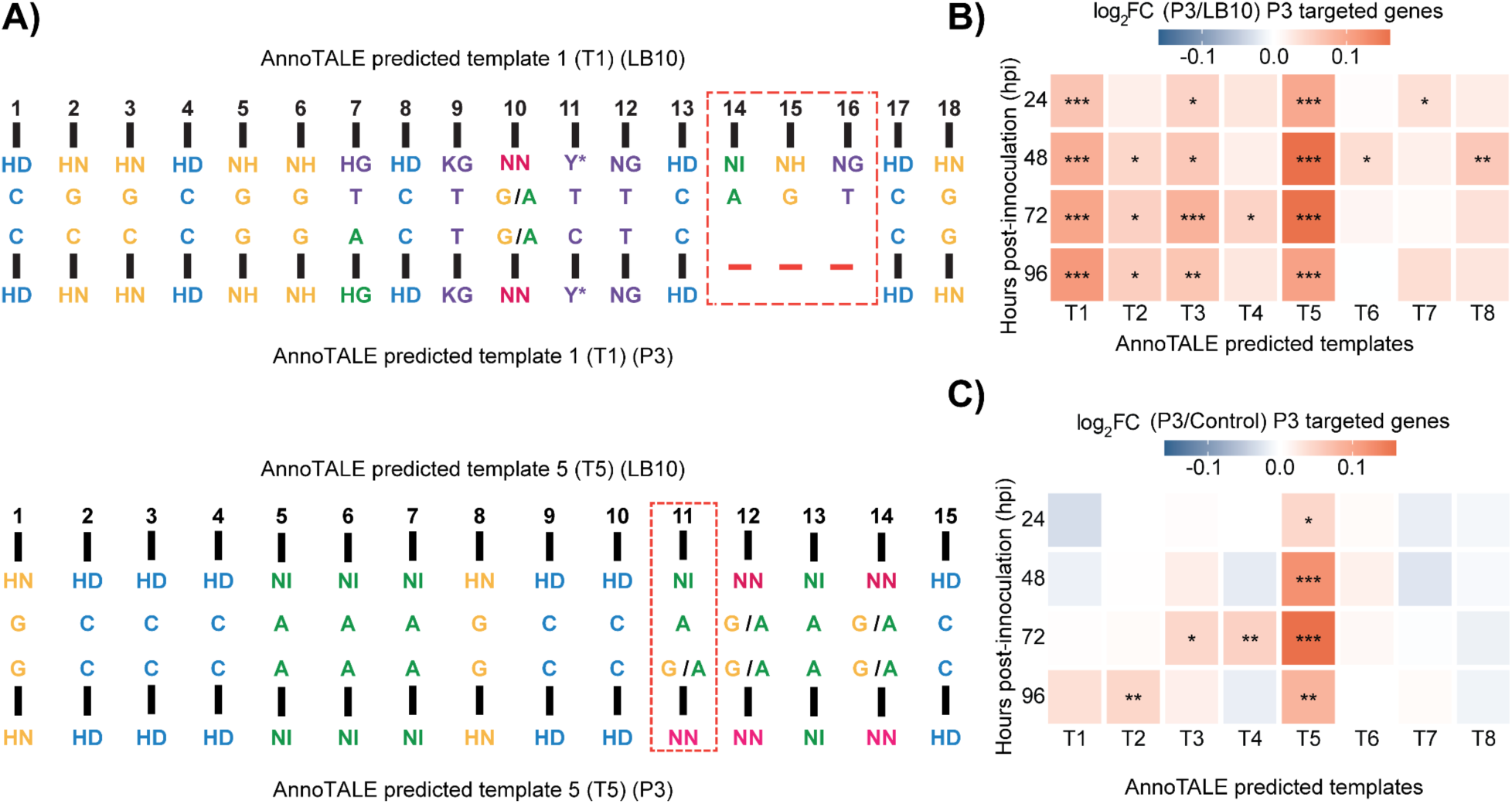
Divergent transcriptional responses of predicted TAL effector targets between LB10 and P3. (A) Alignment of RVDs for TAL1 and TAL5 between LB10 and P3. Differences in RVD composition at specific positions indicate potential variation in EBE recognition specificities between the two strains. Differences in RVD composition between the two strains are highlighted with red dashed boxes; (B) Significance and fold changes in expression between P3 and LB10 for genes predicted to be targeted by P3 TAL effectors across eight TAL templates (T1-T8) at four time points after inoculation; (C) Significance and fold changes in expression between P3 and control for genes predicted to be targeted by P3 TAL effectors across eight TAL templates (T1-T8) at four time points after inoculation.

Given that TAL effector binding to *cis*-regulatory elements could activate gene expression in host genomes (Wang et al., 2017), we compared the expression of genes predicted to be targeted by each P3 TAL effector between P3 and LB10 or control. The 1,000 bp upstream sequences of triticale genes were scanned with each of predicted RVD arrays to identify candidate EBEs. The TALgetter model (Grau et al., 2013) scored each position of the input sequences against each RVD. For each TAL template, the 500 top-scoring EBEs were retained and mapped to the genes whose promoters contained them, yielding about 500 unique candidate target genes per template. Consistent with the greater RVD divergence in template 1, EBE predictions differed more substantially between the two strains, with only 77 commonly predicted genes compared with 196 for template 5. Using the ∼500 predicted target genes by each TAL template in P3, we identified genes that showed significant expression differences between P3 and LB10 across the four time points for template 1, 3 and 5 (Figure 4B). However, only template 5 (TAL5) predicted genes showed significant upregulation under P3 inoculation relative to the control across the four time points (Figure 4C), consistent with the canonical model of TAL effectors as transcriptional activators that induce expression through direct binding to EBEs in target promoters (Moscou and Bogdanove, 2009). To determine whether this expression pattern was specific to P3 TAL5, we performed a reciprocal analysis using genes predicted to be targeted by LB10 TAL5. Although TAL5 differed by only a single RVD between LB10 and P3, the LB10 TAL5-predicted target set showed overall upregulation relative to the control but not relative to P3 (Supplementary Figure 3). We therefore selected P3 TAL5 as a candidate for testing whether this strain-specific transcriptional pattern was associated with a measurable contribution to infection-related phenotypes.

### Temporal phenotypic and transcriptional consequences of TAL5 disruption in P3

To investigate how P3 TAL5 modulates host gene expression, we disrupted and validated the disruption of TAL5 in the P3 background (Supplementary Figure 4). Wild-type P3 and the TAL5 mutant (TAL5KO) were inoculated onto fully extended leaves of triticale “Siskiyou” under the same stage as RNA-seq experiment, to assess their phenotypic responses on virulence. Based on the development of water-soaked symptoms, no obvious difference was observed between the two strains. The visible symptoms produced by the two strains were similar throughout disease development up to 72 hpi (Figure 5A). Because chlorophyll fluorescence imaging was leveraged to detect pathogen-induced physiological changes (Rousseau et al., 2013; Méline et al., 2020), we used this approach to compare phenotypic consequences between TAL5KO- and P3-inoculated leaves that might not be easily captured by RGB images. Maximum quantum efficiency of PSII (F_v_/F_m_) was monitored at five inoculation sites from 8 to 72 hpi, with twelve uniformly developed leaves evaluated per strain at each time point, until water-soaking symptoms became clearly visible (Figure 5A). No significant differences between P3- and TAL5KO-inoculated leaves were detected at 8, 16, 24, or 48 hpi. However, at 72 hpi, TAL5KO-inoculated leaves showed a ∼1.75% reduction in F_v_/F_m_ relative to P3-inoculated leaves when ROIs were summarized using median pixel values (Figure 5B) and a ∼6.2% difference in mean pixel values (Supplementary Figure 5). Because lesion development produced spatially heterogeneous necrotic sectors that could disproportionately influence mean values, we additionally summarized ROIs using median pixel values. This analysis revealed a modest but significant reduction in F_v_/F_m_ in TAL5KO relative to P3 at 72 hpi (paired t-test: p-value = 0.033; ANOVA after controlling for positional and replicate covariates: p-value = 0.029; Figure 5B). Given the small magnitude of this difference and the absence of corresponding changes in visible symptoms, TAL5 disruption appeared to have a modest negative effect on host phenotype performance.

**Figure 5.**
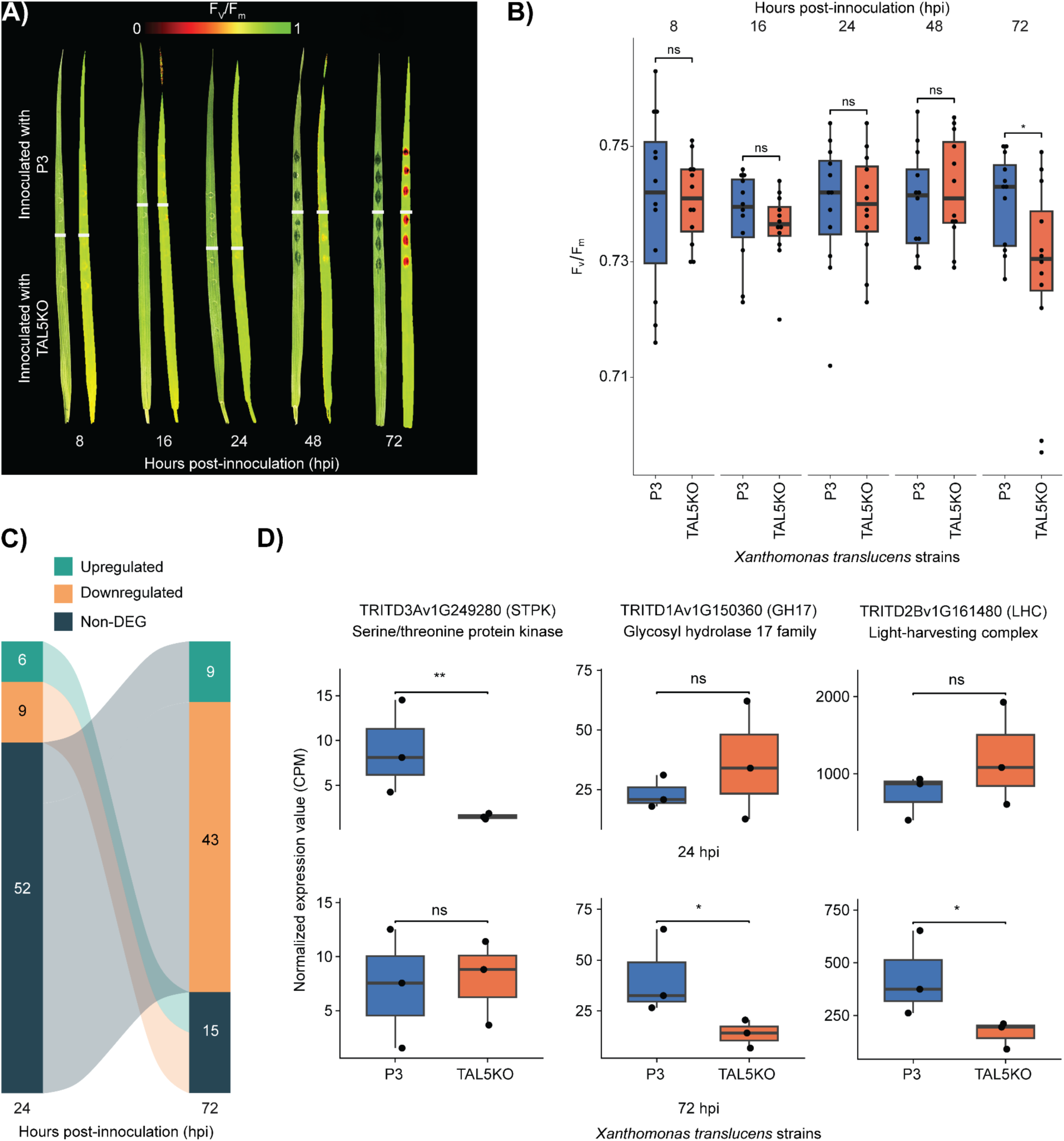
Phenotypic and transcriptional consequences in triticale inoculated with P3 and TAL5KO knockout strains. (A) Temporal disease progression in inoculated leaf regions of triticale infected with P3 or P3-TAL5KO, visualized in RGB and maximum PSII quantum efficiency (F_v_/F_m_) channels at 8, 16, 24, 48 and 72 hours post-inoculation (hpi); (B) Median F_v_/F_m_ values across inoculated regions of interest (ROIs) for P3- and TAL5KO-inoculated leaves at each time point. Total 12 data points for each strain per time point; (C) Numbers of upregulated, downregulated, and non-differentially expressed genes between P3- and TAL5KO-inoculated leaves at 24 and 72 hpi; (D) Normalized expressions in CPM values of three selected genes in P3- and P3-TAL5KO-inoculated leaves at 24 and 72 hpi.

To further examine how transcriptional changes corresponded with phenotypic outcomes, we collected inoculated leaf regions at 24 and 72 hpi from plants grown alongside those used for phenotyping and subjected them to RNA-seq analysis. RNA-seq analysis identified 15 and 52 DEGs between TAL5KO- and P3-inoculated plants at 24 and 72 hpi, respectively, with no DEGs shared between the two time points (Figure 5C; Supplementary Table 6). At both time points, more genes were downregulated than upregulated in TAL5KO relative to P3, with the number of downregulated genes increasing to nearly 5-fold at 72 hpi (Figure 5C), indicating that TAL5 disruption predominantly reduces host gene expression. Of those downregulated DEGs, TRITD3Av1G249280 was specifically downregulated at 24 hpi but not 72 hpi, with almost no expression under TAL5KO inoculated tissue. This gene encodes a serine/threonine protein kinase (STPK), a gene family broadly implicated in plant immune signaling and disease resistance (Afzal et al., 2008) (Figure 5D). At 72 hpi, TRITD1Av1G150360, a glycosyl hydrolase family 17 (GH17) gene, was downregulated in TAL5KO-inoculated tissues compared to P3-inoculated tissues (Figure 5D). GH17 enzymes correspond largely to β-1,3-glucanases (PR-2 family), which degrade β-1,3-glucan in pathogen cell walls and release immunogenic elicitors as part of plant defense (Xu et al., 2016). At 72 hpi, more than half of all DEGs (27 of 52) belonged to photosynthesis-associated functions, including light-harvesting complex (LHC) genes, RuBisCO and ATP synthase subunits. All 27 were downregulated in TAL5KO relative to P3. Among this set, TRITD2Bv1G161480, an ortholog of LHCB6 in *Arabidopsis* and a predicted TAL1 target, was of particular interest given the established role of LHCB6 in PSII supercomplex assembly and its direct link to F_v_/F_m_ variation in maize (Urzinger et al., 2026), offering a plausible mechanistic connection to the reduced F_v_/F_m_ observed in TAL5KO-inoculated leaves at the same time point. Together, these three representative DEGs indicate that TAL5 disruption reduces, rather than derepresses, expression across functionally diverse host genes – defense signaling (STPK), cell wall-associated defense (GH17), and photosynthetic machinery (LHCB6) – consistent with the canonical model of TAL effectors as transcriptional activators, but unexpected in that the induced targets include defense-associated genes rather than genes that promote susceptibility.

## Discussion

During infection by *X. translucens pv. undulosa*, triticale undergoes dynamic phenotypic responses and transcriptional reprogramming shaped by the interplay between host defense and pathogen virulence. In this study, we used time-resolved transcriptomic profiling to characterize the contrasting transcriptional responses of triticale to two *Xtu* strains – the non-virulent strain LB10 and the virulent strain P3 – across four consecutive days of disease progression following inoculation.

Consistent with their contrasting disease outcomes, LB10 and P3 elicited distinct temporal transcriptional responses. This divergence was most pronounced at 24 hpi, when approximately twice as many DEGs were detected in response to LB10 as to P3, a pattern in line with the rapid induction of defense-related genes reported in incompatible interactions (Wang et al., 2021). Specifically, GO terms uniquely enriched in LB10 pointed toward defense-associated processes. For example, monoterpene biosynthetic process (GO:0043693) was specifically enriched for LB10, a process linked to resistance against *Xanthomonas* pathogens in rice (Taniguchi et al., 2014). In the compatible response induced by P3, the lower number of DEGs at earlier time points is likely due to the suppression of plant defense responses by pathogen effectors. The overall transcriptomic pattern was dominated by photosynthesis-associated changes (GO:0009521 “photosystem components”) and primary-metabolism-associated changes (GO:0006520 “amino acid metabolism”) that intensified as symptoms developed. This transcriptional response, centered on photosynthetic and metabolic functions rather than defense signaling, more likely reflects the physiological consequences of advancing disease than an effective defense response in P3.

As a hexaploid species, triticale integrates transcriptional inputs from three subgenomes, and their coordination shapes phenotypic outcomes. Each syntenic triad was assigned to one of seven categories according to the relative expression of its A, B, and R homoelogs (Figure 2A). Under the null expectation, the three subgenomes should contribute equally to dominant and suppressed homoelogs – as largely observed in bread wheat (Ramírez-González et al., 2018). In triticale, however, R-derived homoelogs were 3x overrepresented in the suppressed state relative to A- and B-derived homoelogs regardless of conditions or time points, whereas no such bias was observed in the dominant state. A precedent for such genome-specific bias exists in wheat-rye hybrids, where rye rDNA loci are silenced through nucleolar dominance (Neves et al., 1997). Although restricted to rDNA, this illustrates how parental genomes can interact unequally in newly formed allopolyploids. Unlike the largely balanced homoelog expression of natural hexaploid wheat like bread wheat, the recent artificial origin of triticale entails incomplete regulatory compatibility between the rye genome and the wheat-derived regulatory environment, resulting in preferential repression of R homoelogs. This asymmetry makes the R subgenome particularly interesting in the context of BLS, as the only major resistance gene mapped to date in triticale – a dominant locus effective against strain LB10 – resides on chromosome 5R (Wen et al., 2018), and its function was partially inferred to be modulated by poor expression of rye genes, consistent with the suppression pattern we observed.

Beyond the regulatory architecture of the host genome, susceptibility to *Xanthomonas* is frequently determined by the ability of pathogens to co-opt host transport machinery. In the rice-*X. oryzae pv. oryzae* pathosystem, TAL effectors bind the promoters of SWEET sucrose transporter genes and induce their expression, releasing sugars into the apoplast to support bacterial multiplication (Streubel et al., 2013). In other cereals, TAL effector targets are not restricted to sugar transporters. In rice, Tal2g of *X. oryzae* pv. *oryzicola* induces the sulfate transporter gene *OsSULTR3;6* (Cernadas et al., 2014). In wheat, TAL8 of *X. translucens* pv. *undulosa* induces *TaNCED-5BS*, which encodes the rate-limiting enzyme of abscisic acid biosynthesis and promotes susceptibility by altering host water status (Peng et al., 2019). These different observations suggest that TAL effector targets may diverge across *Xanthomonas*-host combinations rather than converging on a single class of susceptibility gene. Because our clusters were defined by temporal expression trajectories across both interactions rather than by pairwise contrasts at individual time points, they resolved temporally structured processes that single-time point comparisons could not. Of 18 SWEET genes classified as DEGs in either LB10- or P3-inoculated samples across four time points, only two – TRITD3Av1G236060 and TRITD5Bv1G016340 – showed upregulation, but neither was directly targeted by any TAL effector template. However, cluster 4 showed a sharp late-stage induction in P3 that exceeded the more modest increase observed in LB10 and was enriched for several sugar transport functions, including sugar transmembrane transporter activity (GO:0051119), sucrose transport (GO:0015770), and D-glucose import (GO:0046323). The enrichment of these transport functions suggest that susceptibility might involve activation of alternative host sugar transport pathways beyond the SWEET-mediated mechanisms well characterized in the *rice-Xanthomonas* pathosystem.

To understand how TAL targeted genes responded, we used AnnoTALE to predict EBEs for the TAL effector templates encoded by LB10 and P3. Of the eight templates, only TAL1 and TAL5 differed in RVD compositions between the two strains. Interestingly, only genes predicted as TAL5 targets were expressed at higher levels in P3 than in both LB10 and mock-inoculated plants across four time points (Figure 4B and 4C), suggesting TAL5 may contribute in strain-specific transcriptional effects. Subsequently, inoculation with TAL5KO strain in triticale yielded more downregulated than upregulated genes relative to wild-type P3 at both 24 and 72 hpi, consistent with the established mechanism of TAL effectors as transcriptional activators (Bogdanove et al., 2010; Wang et al., 2017). However, an important caveat was that none of the predicted TAL5 target genes were themselves differentially expressed in the knockout comparison, suggesting that the observed changes might likely be indirect or target predictions were not accurate. Although, in a recent study in *Xanthomonas campestris pv. Campestris* (Xcc)-*Brassica* interaction demonstrated that Xcc pathogenicity in *Brassica* relies predominantly on non-TAL Xop effectors that can work for both disease suppression and progression depending on the host genotype (Chen et al. 2026). One differentially expressed gene, TRITD2Bv1G161480, did carry a predicted EBE, but for TAL1 rather than TAL5. Expression of this gene was reduced in P3 relative to the control from 48 hpi onward, consistent with the susceptible symptoms observed in four time points experiment. In the TAL5KO/P3 comparison, the same gene was further reduced in TAL5KO relative to P3 at 72 hpi, paralleled by lower F_v_/F_m_ in TAL5KO-than in P3-inoculated leaves. Across both experiments, this gene and sixteen additional LHC genes declined in the same rank order as symptom severity at 72 hpi (Supplementary Table 6), a pattern consistent with the general cost of biotic stress to host photosynthetic capacity (Bilgin et al., 2010). Together, our findings suggest that the effects of TAL effectors on host transcriptional reprogramming may vary across TAL effector templates and different host-*Xanthomonas* interaction systems.

## Materials and Methods

### Plant growth conditions and bacteria strain inoculations

The triticale cultivar Siskiyou (CI 17603) was developed by the International Maize and Wheat Improvement Center (CIMMYT) in collaboration with the University of California, Davis, and released in California in 1978 (Qualset et al., 1978). Two *Xanthomonas translucens pv. undulosa* strains, LB10 and P3, originally collected in North Dakota, were used for bacterial infiltration (Adhikari et al., 2011). Siskiyou was previously shown to exhibit differential responses to LB10 and P3 (Sapkota et al., 2018; Wen et al., 2018). For RNA-seq sample collection, plants were grown in cones (4 × 13 cm), with two plants per cone. Cones were filled with Sunshine LC1 growing mix (Sun Gro Horticulture Distribution Inc.), and a small amount of Osmocote 15-9-12 fertilizer (Everris NA Inc.) was added to each cone. Plants were grown in a greenhouse environment with day/night temperatures of 24°C/18°C under a 14-h photoperiod, with average humidity maintained at 40-50%.

### Inoculum cultivation and inoculation

The inoculum was prepared according to the protocol (Adhikari et al., 2011) where both bacterial strains LB10 and P3 were streaked from the stock culture in −80°C on Wilbrink’s agar (WBA) plate. The plates were incubated at 28°C for two days. The bacterial inoculum was prepared by washing bacterial culture in a 1x PBS buffer (saline buffer; 800mg NaCl, 200mg KCl, 144mg Na_2_HPO_4_, 245mg KH_2_PO_4_ in 1 L of distilled water, pH = 7.4). The final concentration of the inoculum was adjusted to optical density of 0.2 at 600 nm (OD_600_) using a spectrophotometer. The inoculations were done at the three-leaf seedling stage (14 days after planting) using a spot-infiltration method with a 1-mL needleless syringe as previously described (Sapkota et al., 2018). The mock inoculation was done by using 1x PBS buffer alone.

### RNA-seq data generations

Tissue samples were collected at 24, 48, 72, and 96 hours post-infiltration. Briefly, a leaf section about two inches with four infiltration spots was cut using sterilized scissors and placed into a 2 ml micro-centrifuge tube. Tissue samples were collected from three biological replicates for each strain at each time point. The samples were immediately transferred to liquid nitrogen and stored at −80°C. The total RNA was extracted using the RNeasy mini kit (Qiagen). RNA concentration was then measured using the NanoDrop One/OneC UV-Vis spectrophotometer (Thermo Fisher Scientific Inc., USA). Quality control was assessed through multiple criteria including RNA quantification, agarose gel electrophoresis, RNA Quality Number/RNA Integrity Number (RQN/RIN) analysis, 260/280 and 260/230 absorbance ratios. The mRNA enrichment, library preparation and sequencing were done by Novogene Inc. USA (Sacramento, CA). Non-stranded RNA-Seq libraries were prepared and sequenced on Illumina Novaseq platforms using a 150bp paired-end configuration.

### RNA-seq data processing

Raw RNA-seq reads were preprocessed using fastp v1.0.1 (Chen et al., 2018) to remove adapter sequences and low-quality reads. Reference genome sequences, cDNA sequences, and genome annotations for durum wheat (Svevo v1) were obtained from Ensembl Plants release 59, while the corresponding resources for rye (Lo7 v3) were obtained from the e!DAL Plant Genomics & Phenomics Research Data Repository (https://doi.org/10.5447/IPK/2020/33). Reference cDNA sequences from the durum wheat and rye genomes were concatenated to construct a combined reference transcriptome for triticale. A transcriptome index was built using kallisto v0.51.1 (Bray et al., 2016), and preprocessed reads were aligned to the triticale transcriptome using default parameters. Kallisto was used to estimate transcript-level abundance.

A transcript database was constructed from the reference genome annotations using the makeTxDbFromGFF function implemented in the Bioconductor package txdbmaker v1.4.2. Transcript-to-gene relationships were established using a custom annotation table containing corresponding transcript and gene identifiers. Transcript-level abundance estimates were summarized to the gene level using the R package tximport v1.36.1 (Soneson et al., 2015), with countsFromAbundance = "lengthScaledTPM" to generate gene-level count matrices while accounting for transcript length. Transcripts without detected reads across all samples were excluded prior to downstream analysis.

The resulting gene-level count matrix was imported into DESeq2 v1.48.2 (Love et al., 2014; Putri et al., 2022) for differential expression analysis. For the time-course experiment, LB10- and P3-inoculated samples were independently compared with their corresponding control samples at each time point. For the validation experiment, P3 TAL5 knockout mutant-inoculated samples were compared with samples inoculated with P3. Genes with a false discovery rate (FDR)-adjusted p-value < 0.05 and an absolute log2fold-change ≥ 1 were considered DEGs.

### Identification of syntenic homoeolog triads across the A, B, and R subgenomes

Hardmasked genomes and associated annotation files of durum wheat and rye were downloaded from Ensembl Plants 59. SynMap in CoGe (https://genomevolution.org/coge/) was implemented to generate synteny maps and syntenic gene pairs. Specifically, the LAST alignment algorithm (Kiełbasa et al., 2011) was implemented to identify coding sequence similarities. DAGChainer options were set as -D 20 and -A 5. Syntenic blocks were merged by Quota Align Merge (-Dm 20). The ratio of coverage depth was set as 1:2 for rye relative to durum wheat. We generated two lists of syntenic gene pairs --within durum wheat, between durum wheat and rye. Because durum wheat is allotetraploid (AABB), we considered genes reciprocally identified as syntenic between chromosomes in durum wheat as the final syntenic gene list in durum wheat. In addition, any rye genes that were not identified as syntenic genes relative to durum wheat were considered as singletons. To generate unique 1:1:1 syntenic gene triplets among the durum wheat A and B subgenomes and rye, each rye gene was required to be syntenic with both members of a corresponding durum wheat homoelogous gene pair. Gene pairs located on non-homologous chromosomes were excluded in the analysis.

### Relative expression patterns of syntenic homoeologs across A:B:R triads in triticale

Relative homoeolog expression was evaluated following the general approach described by (Ramírez-González et al., 2018). To exclude syntenic triads with low expression, only triads with the summed expression of its A, B, and R homoeologs exceeding 0.5 counts per million (CPM) were considered in the analysis. We further required each triad to have at least one homoeolog with CPM >0.5. As there was a slight variation in triad expression between time point because of the dynamic temporal shifts in expression, we consolidated triads from 24, 48, 72, and 96 for each treatment resulting in a final set of 8,493 expressed triads for control, 8,684 expressed triads for LB10 and 8,556 triads for P3 retained for relative expression analysis.

To quantify the relative contribution of each homoeolog to the total expression of a triad, CPM values were normalized within each triad as follows:

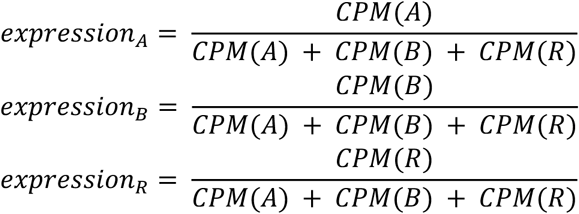

where CPM(A), CPM(B), and CPM(R) represent the expression levels of the A-, B-, and R-subgenome homoeologs, respectively. Thus, the relative contributions of the three homoeologs within each triad sum to 1. Based on the relative contributions of the A, B, and R homoeologs, each triad was assigned to one of seven homoeolog expression-bias categories: A-dominant, B-dominant, R-dominant, A-suppressed, B-suppressed, R-suppressed, or balanced. Specifically, a triad was classified as A-, B-, or R-dominant when the relative contribution of the corresponding homoeolog was ≥0.8. Conversely, a triad was classified as A-, B-, or R-suppressed when the relative contribution of the corresponding homoeolog was ≤0.2. Triads that did not meet the criteria for either dominant or suppressed expression were classified as balanced. Relative subgenome contributions and homoeolog expression-bias categories were visualized using ternary plots generated with the R package ggtern v4.0.0 (Hamilton and Ferry, 2018). This relative homoeolog expression analysis was separately performed for each condition (control, LB10 and P3) at each time point (24, 48, 72, and 96 hpi).

### Temporal expression pattern clustering

Across the four time points (24, 48, 72, and 96 hpi), a total of 15,197 unique DEGs were identified from eight differential expression comparisons, including LB10 versus control and P3 versus control at each time point. Log_2_fold-change values for each gene were standardized across the eight datasets using z-score normalization, resulting in a mean of 0 and a standard deviation of 1 for each gene. The resulting normalized expression matrix was analyzed using kmeans() in R to identify clusters with distinct regulation patterns. To determine the optimal number of clusters (k), the within-cluster sum of squares (WSS) was calculated for k values ranging from 1 to 20. The second derivative of the WSS curve was then computed to evaluate changes in the rate of WSS reduction. The optimal k value was determined as 16 based on the rate of WSS reduction and the inflection point identified through the second derivative of the WSS curve (Supplementary Figure 2).

### GO database construction

To improve Gene Ontology (GO) annotations for triticale, GOMAP (Gene Ontology Meta Annotator for Plants) (Wimalanathan and Lawrence-Dill, 2021) was used to assign GO terms to genes based on the protein sequences of durum wheat and rye. The resulting GO annotations were uploaded to https://github.com/zlianglab/Triticale_BLS.

### Gene ontology enrichment analysis

GO enrichment analysis was performed for each gene set using GOATOOLS v1.5.1 (Grau et al., 2016; Klopfenstein et al., 2018). The background comprised 70,131 genes with a mean CPM > 1 across all time points. P-values were adjusted using Bonferroni correction and GO terms with adjusted p < 0.05 were considered significantly enriched.

To reduce redundancy for GO enrichment in each functional cluster, 1,961 enriched GO terms from the 16 clusters were consolidated into 1,080 unique terms and analyzed using REVIGO (Supek et al., 2011). REVIGO was run with medium redundancy reduction (0.7), the SimRel similarity measure, *Arabidopsis thaliana* as the reference species. Semantically related terms were grouped, and the most frequent term in each group was selected as its representative. After excluding cellular-component and low-frequency categories, 27 representative GO terms were assigned to the 16 clusters, min-max normalized, and visualized using pheatmap v1.0.13.

### Coding sequence and upstream region similarity among homoeologs within a triad

Of three homoeologs in a triad, the CDS sequences along with 1,000-bp upstream sequences upstream from the start codon were extracted using pyfaidx v.0.9.0.4 (Shirley et al., 2015). Afterwards, pairwise sequence similarity scores of each of the triad classes were calculated using mafft v7.525 (Katoh and Standley, 2013). The average of the three pairwise similarity scores within each triad was calculated to represent the overall CDS or upstream sequence similarity.

### Motif enrichment analysis

To identify enriched sequence motifs associated with each gene cluster, 1,000-bp sequences upstream of the start codon were extracted for genes within each cluster using the Python module pyfaidx v0.9.0.4 (Shirley et al., 2015). The start codon was used as the reference because transcription start sites were not consistently annotated in the reference genomes. *De novo* motif discovery was performed using STREME using 1,000bp-sequences of all expressed genes as negative control, and identified motifs were compared with known TF binding motifs using TOMTOM, both implemented in MEME Suite v5.5.8 (Bailey et al., 2015). For each cluster, upstream sequences of genes within the cluster were used for motif discovery, while upstream sequences from 70,131 expressed genes were used as the background to identify cluster-enriched motifs. STREME searches were restricted to motifs ranging from 8 to 20 nucleotides in width. To infer potential TFs associated with the identified motifs, STREME-derived motifs were compared against known TF-binding motifs from PlantTFDB v5.0 (Jin et al., 2016). Identified motifs were compared with known transcription factor binding motifs from *Arabidopsis thaliana* in PlantTFDB v5.0 (Jin et al., 2016) using TOMTOM with a significance threshold of 0.05.

### Identification of SWEET genes in Triticale

SWEET genes in triticale were identified based on the presence of the conserved MtN3/saliva domain (Pfam accession PF03083). The corresponding hidden Markov model (HMM) profile was retrieved from the Pfam-A database using hmmfetch implemented in HMMER v3.4 (Finn et al., 2011). The extracted HMM profile was then used to search all triticale protein sequences using hmmsearch with default parameters to identify candidate SWEET genes.

### TAL effector binding sequence predictions

Genome sequences of P3 (Peng et al., 2019) and LB10 (unpublished) were used for TAL effector analysis. The start codon was used as the reference position because transcription start sites (TSSs) are incompletely annotated in these genomes. To identify potential TAL effector binding sites for each host gene, 1,000-bp sequences upstream of the TSS were extracted from the corresponding durum wheat or rye genome assemblies using the pyfaidx Python module (Shirley et al., 2015). TAL effector binding sites and RVD arrays of predicted TAL effectors were identified using AnnoTALE v1.5 (Erkes et al., 2017).

### TAL5 disruption and functional characterization

The gene disruption for TAL5 in P3 followed a similar procedure as previously reported (Peng et al., 2019). Briefly, a highly conserved region in the N terminus of TALE genes was amplified and cloned into the suicide vector pKnock-Km. The resulting plasmid was transformed into P3 through bacterial conjugation. The mutants were screened with TAL5 gene specific primers (Supplementary Table 7) as described in a prior study (Peng et al., 2019) and confirmed by the whole genome sequencing performed by Plasmidsaurus (Louisville, KY). To test the regulation by TAL5 in triticale, the wild type P3 and this mutant were infiltrated onto Sysikiou and leaf tissues were collected at 8 hpi, 16 hpi, 24 hpi, 48 hpi and 72 hpi. Inoculated leaf tissues from P3 and the TAL5 knockout strains were collected at 24 hpi and 72 hpi in three biological replicates for RNA-seq analysis.

### Chlorophyll fluorescence imaging of plants inoculated with P3 and TAL5 knockout strains

For phenotyping, the most recently fully expanded leaves of 14-day-old plants were inoculated with both P3 and TAL5KO at separate sites in the center region of each leaf. Twelve leaves were evaluated at each time point (8, 16, 24, 48, and 72 hpi). For 24 and 72 hpi, leaf samples for imaging were collected in tandem with the RNAseq samples. To control positional effects, P3 and TAL5KO were assigned to opposite inoculation sites on six leaves, with their positions reversed on the remaining six leaves. Chlorophyll fluorescence imaging was conducted using a PlantExplorer Pro+ system (PhenoVation B.V., Netherlands), following a previously employed protocol (Fernandez et al., 2026). Leaves were dark-adapted for 30 min prior to imaging. Minimal (F_0_) and maximal (Fm) fluorescence were measured using a saturating light pulse, and the maximum quantum efficiency of PSII photochemistry was calculated as F_v_/F_m_ = (F_m_ − F_0_)/F_m_. At each time point, three inoculation spots were selected in one ROI. Within each ROI, mean and median F_v_/F_m_ value was calculated using CropReporter™ v5.9.0-64b software. Paired t-tests were used to compare F_v_/F_m_ between P3-and TAL5KO-inoculated sites. To account for potential positional effects, an additional ANOVA was performed with strain and inoculation position included as fixed effects.

## Author contributions

Z.H.L. conceived the experiments. Z.K.L. designed the analytical framework. F.H., F.M., E.C.F., and S.W. collected the data. F.H. and Z.K.L. performed the data analyses, with B.D. and C.Y. contributing to discussions and development of the initial analytical approaches. G.S. and Z.H.L. performed strain knockout experiments. F.H., F.M., E.C.F., Z.H.L., and Z.K.L. wrote the manuscript. All authors reviewed and approved the final manuscript.

## Data Availability

RNA-seq data from mock-, LB10-, and P3-inoculated leaf regions collected at 24, 48, 72, and 96 hours post-inoculation, together with RNA-seq data from P3- and TAL5KO-inoculated leaf collected at 24 and 72 hours post-inoculation, with three biological replicates per treatment at each time point, have been deposited in the NCBI Sequence Read Archive (SRA) under BioProject accession PRJNA1522063. All custom scripts used for data processing, statistical analysis, and visualization in this study are publicly available on GitHub at https://github.com/zlianglab/Triticale_BLS.

## Supplementary Information

### Supplementary Figures

**Supplementary Figure S1.** Dynamic shifts in triad categories across treatments and time points. (A) Shifts between LB10 and P3 at each of the four time points; temporal shifts across the four time points under (B) control and (C) LB10 conditions.

**Supplementary Figure S2.** The Elbow method for selecting the optimal number of gene clusters.

**Supplementary Figure S3.** Significance and fold changes in expression (A) between LB10 and P3, and (B) between LB10 and control. Genes predicted to be targeted by LB10 TAL effectors across eight TAL templates (T1-T8) were used for comparison at each of four post-inoculation time points.

**Supplementary Figure S4.** The scheme and confirmation of TAL5 knockout strain generation. A construct based on pKnock-Km (suicide vector) was made by adding a 500 bp fragment of TALE5 (N-terminus) and transformed into *Xanthomonas tranlucens pv. undolusa* strain P3 where the construct was integrated into the bacterial genome based on homologous recombination.

**Supplementary Figure S5.** Mean F_v_/F_m_ values for inoculated regions of interest (ROIs) between P3-and TAL5KO-inoculated leaves at each time point.

### Supplementary Tables

**Supplementary Table 1.** Uniquely enriched GO terms in LB10- and P3-inoculated samples across four time points.

**Supplementary Table 2.** Enriched GO terms in R genes for R-suppressed triads.

**Supplementary Table 3.** Original and consolidated GO terms across sixteen gene clusters.

**Supplementary Table 4.** Identified TF binding motifs across gene clusters.

**Supplementary Table 5.** The list of SWEET genes identified in Triticale.

**Supplementary Table 6.** Differentially expressed genes identified between TAL5KO and P3 inoculated samples.

**Supplementary Table 7.** Primer sequences used in TAL5 knockout experiments.

## Acknowledgements

This work was supported by the Agricultural Research Service Grants #58-3060-3-016 and #58-3060-3-022 from the U.S. Department of Agriculture (USDA), and the North Dakota State University start-up fund to Z.K.L. This research was also supported by North Dakota Wheat Commission and the National Institute of Food and Agriculture, USDA, under Hatch project number ND02248 awarded to Z.H.L. The findings and conclusions in this preliminary publication have not been formally disseminated by the U. S. Department of Agriculture and should not be construed to represent any agency determination or policy. This work, using resources of the Center for Computationally Assisted Science and Technology (CCAST) at North Dakota State University, was made possible in part by the National Science Foundation Major Research Instrumentation (MRI) Award No. 2019077.

## Notes

### Competing Interest Statement

The authors have declared no competing interest.

## References

Acharya K, Liu Z, Schachterle J, Kumari P, Manan F, Xu SS, Green AJ, Faris JD (2024) Genetic mapping of QTLs for resistance to bacterial leaf streak in hexaploid wheat. Theor Appl Genet 137: 265

Adhikari TB, Hansen JM, Gurung S, Bonman JM (2011) Identification of New Sources of Resistance in Winter Wheat to Multiple Strains of Xanthomonas translucens pv. undulosa. Plant Dis 95: 582–588

Afzal AJ, Wood AJ, Lightfoot DA (2008) Plant receptor-like serine threonine kinases: roles in signaling and plant defense. Mol Plant Microbe Interact 21: 507–517

Bailey TL, Johnson J, Grant CE, Noble WS (2015) The MEME Suite. Nucleic Acids Res 43: W39–W49

Bilgin DD, Zavala JA, Zhu J, Clough SJ, Ort DR, DeLucia EH (2010) Biotic stress globally downregulates photosynthesis genes. Plant Cell Environ 33: 1597–1613

Bird KA, VanBuren R, Puzey JR, Edger PP (2018) The causes and consequences of subgenome dominance in hybrids and recent polyploids. New Phytol 220: 87–93

Bogdanove AJ, Schornack S, Lahaye T (2010) TAL effectors: finding plant genes for disease and defense. Curr Opin Plant Biol 13: 394–401

Bragard C, Singer E, Alizadeh A, Vauterin L, Maraite H, Swings J (1997) Xanthomonas translucens from Small Grains: Diversity and Phytopathological Relevance. Phytopathology 87: 1111–1117

Bray NL, Pimentel H, Melsted P, Pachter L (2016) Erratum: Near-optimal probabilistic RNA-seq quantification. Nat Biotechnol 34: 888

Castro-Moretti FR, Gentzel IN, Mackey D, Alonso AP (2020) Metabolomics as an Emerging Tool for the Study of Plant-Pathogen Interactions. Metabolites. doi: 10.3390/metabo10020052

Cernadas RA, Doyle EL, Niño-Liu DO, Wilkins KE, Bancroft T, Wang L, Schmidt CL, Caldo R, Yang B, White FF, et al (2014) Code-assisted discovery of TAL effector targets in bacterial leaf streak of rice reveals contrast with bacterial blight and a novel susceptibility gene. PLoS Pathog 10: e1003972

Chen S, Zhou Y, Chen Y, Gu J (2018) fastp: an ultra-fast all-in-one FASTQ preprocessor. Bioinformatics 34: i884–i890

Dong S, Adams KL (2011) Differential contributions to the transcriptome of duplicated genes in response to abiotic stresses in natural and synthetic polyploids. New Phytol 190: 1045– 1057

Erkes A, Mücke S, Reschke M, Boch J, Grau J (2019) PrediTALE: A novel model learned from quantitative data allows for new perspectives on TALE targeting. PLoS Comput Biol 15: e1007206

Erkes A, Reschke M, Boch J, Grau J (2017) Evolution of Transcription Activator-Like Effectors in Xanthomonas oryzae. Genome Biol Evol 9: 1599–1615

Fernandez EC, Tu G, Dai W, Yang S, Liu Z, Grzybowski M, Liang Z (2026) Spatial coordination between leaf gradient and temperature response in barley. Plant J 126: e70857

Finn RD, Clements J, Eddy SR (2011) HMMER web server: interactive sequence similarity searching. Nucleic Acids Res 39: W29–37

Garcia-Seco D, Chiapello M, Bracale M, Pesce C, Bagnaresi P, Dubois E, Moulin L, Vannini C, Koebnik R (2017) Transcriptome and proteome analysis reveal new insight into proximal and distal responses of wheat to foliar infection by Xanthomonas translucens. Sci Rep 7: 10157

Grau J, Reschke M, Erkes A, Streubel J, Morgan RD, Wilson GG, Koebnik R, Boch J (2016) AnnoTALE: bioinformatics tools for identification, annotation, and nomenclature of TALEs from Xanthomonas genomic sequences. Sci Rep 6: 21077

Grau J, Wolf A, Reschke M, Bonas U, Posch S, Boch J (2013) Computational predictions provide insights into the biology of TAL effector target sites. PLoS Comput Biol 9: e1002962

Gutierrez-Castillo DE, Barrett E, Roberts R (2024) A recently collected Xanthomonas translucens isolate encodes TAL effectors distinct from older, less virulent isolates. Microb Genom. doi: 10.1099/mgen.0.001177

Hamilton NE, Ferry M (2018) ggtern: Ternary Diagrams Using ggplot2. J Stat Soft 87: 1–17

He F, Wang W, Rutter WB, Jordan KW, Ren J, Taagen E, DeWitt N, Sehgal D, Sukumaran S, Dreisigacker S, et al (2022) Genomic variants affecting homoeologous gene expression dosage contribute to agronomic trait variation in allopolyploid wheat. Nat Commun 13: 826

Jin J, Tian F, Yang D-C, Meng Y-Q, Kong L, Luo J, Gao G (2016) PlantTFDB 4.0: toward a central hub for transcription factors and regulatory interactions in plants. Nucleic Acids Res 45: D1040–D1045

Ji Z, Ji C, Liu B, Zou L, Chen G, Yang B (2016) Interfering TAL effectors of Xanthomonas oryzae neutralize R-gene-mediated plant disease resistance. Nature Communications 7: 13435

de Jong GW, Adams KL (2023) Subgenome-dominant expression and alternative splicing in response to Sclerotinia infection in polyploid Brassica napus and progenitors. Plant J 114: 142–158

Katoh K, Standley DM (2013) MAFFT multiple sequence alignment software version 7: improvements in performance and usability. Mol Biol Evol 30: 772–780

Kiełbasa SM, Wan R, Sato K, Horton P, Frith MC (2011) Adaptive seeds tame genomic sequence comparison. Genome Res 21: 487–493

Klopfenstein DV, Zhang L, Pedersen BS, Ramírez F, Warwick Vesztrocy A, Naldi A, Mungall CJ, Yunes JM, Botvinnik O, Weigel M, et al (2018) GOATOOLS: A Python library for Gene Ontology analyses. Sci Rep 8: 10872

Ledman KE, Osdaghi E, Curland RD, Liu Z, Dill-Macky R (2023) Epidemiology, host resistance, and genomics of the small grain cereals pathogen Xanthomonas translucens: New advances and future prospects. Phytopathology 113: 2037–2047

Lee JS, Adams KL (2020) Global insights into duplicated gene expression and alternative splicing in polyploid Brassica napus under heat, cold, and drought stress. Plant Genome 13: e20057

van Loon LC, Rep M, Pieterse CMJ (2006) Significance of inducible defense-related proteins in infected plants. Annu Rev Phytopathol 44: 135–162

Love MI, Huber W, Anders S (2014) Moderated estimation of fold change and dispersion for RNA-seq data with DESeq2. Genome Biol 15: 550

Ma X-F, Gustafson JP (2008) Allopolyploidization-accommodated genomic sequence changes in triticale. Ann Bot 101: 825–832

Méline V, Brin C, Lebreton G, Ledroit L, Sochard D, Hunault G, Boureau T, Belin E (2020) A Computation Method Based on the Combination of Chlorophyll Fluorescence Parameters to Improve the Discrimination of Visually Similar Phenotypes Induced by Bacterial Virulence Factors. Front Plant Sci 11: 213

Moscou MJ, Bogdanove AJ (2009) A simple cipher governs DNA recognition by TAL effectors. Science 326: 1501

Murcia Bermudez JM, Gonzalez-Bello DA, Ramirez R, Poudel-Ward B (2026) First Report of Xanthomonas translucens pv. undulosa Causing Bacterial Leaf Streak on Wheat in Arizona, U.S.A. Plant Disease. doi: 10.1094/PDIS-11-25-2199-PDN

Neves N, Silva M, Heslop-Harrison JS, Viegas W (1997) Nucleolar dominance in triticales: control by unlinked genes. Chromosome Res 5: 125–131

Ngou BPM, Ahn H-K, Ding P, Jones JDG (2021) Mutual potentiation of plant immunity by cell-surface and intracellular receptors. Nature 592: 110–115

Ohme-Takagi M, Shinshi H (1995) Ethylene-inducible DNA binding proteins that interact with an ethylene-responsive element. Plant Cell 7: 173–182

Oliva R, Ji C, Atienza-Grande G, Huguet-Tapia JC, Perez-Quintero A, Li T, Eom J-S, Li C, Nguyen H, Liu B, et al (2019) Broad-spectrum resistance to bacterial blight in rice using genome editing. Nat Biotechnol 37: 1344–1350

Osdaghi E, Taghavi SM, Aliabadi AA, Khojasteh M, Abachi H, Moallem M, Mohammadikhah S, Shah SMA, Chen G, Liu Z (2023) Detection and Diagnosis of Bacterial Leaf Streak on Small Grain Cereals: From Laboratory to Field. Phytopathology 113: 2024–2036

Peng J, Liu S, Wu J, Liu T, Liu B, Xiong Y, Zhao J, You M, Lei X, Ma X (2024) Genome-wide analysis of the oat (Avena sativa) HSP90 gene family reveals its identification, evolution, and response to abiotic stress. Int J Mol Sci 25: 2305

Peng Z, Huguet-Tapia JC, White F (2019a) *Xanthomonas translucens* pv. *undulosa* strain P3 chromosome, complete genome. Xanthomonas translucens commandeers the host rate-limiting step in ABA biosynthesis for disease susceptibility

Peng Z, Hu Y, Zhang J, Huguet-Tapia JC, Block AK, Park S, Sapkota S, Liu Z, Liu S, White FF (2019b) Xanthomonas translucens commandeers the host rate-limiting step in ABA biosynthesis for disease susceptibility. Proc Natl Acad Sci U S A 116: 20938–20946

Powell JJ, Fitzgerald TL, Stiller J, Berkman PJ, Gardiner DM, Manners JM, Henry RJ, Kazan K (2017) The defence-associated transcriptome of hexaploid wheat displays homoeolog expression and induction bias. Plant Biotechnol J 15: 533–543

Putri GH, Anders S, Pyl PT, Pimanda JE, Zanini F (2022) Analysing high-throughput sequencing data in Python with HTSeq 2.0. Bioinformatics 38: 2943–2945

Qualset CO, Vogt HE, Gustafson JP, Zilinsky FJ, Prato JD, Beatty KD (1978) Siskiyou - a triticale variety for northern California. Calif Agric (Berkeley) 4, 5

Raffaele S, Rivas S (2013) Regulate and be regulated: integration of defense and other signals by the AtMYB30 transcription factor. Front Plant Sci 4: 98

Ramírez-González RH, Borrill P, Lang D, Harrington SA, Brinton J, Venturini L, Davey M, Jacobs J, van Ex F, Pasha A, et al (2018) The transcriptional landscape of polyploid wheat. Science. doi: 10.1126/science.aar6089

Rezzonico F, Rupp O, Fahrentrapp J (2017) Pathogen recognition in compatible plant-microbe interactions. Sci Rep 7: 6383

Roman-Reyna V, Heiden N, Butchacas J, Toth H, Cooperstone JL, Jacobs JM (2024) The timing of bacterial mesophyll infection shapes the leaf chemical landscape. Microbiology Spectrum. doi: 10.1128/spectrum.04138-23

Rousseau C, Belin E, Bove E, Rousseau D, Fabre F, Berruyer R, Guillaumès J, Manceau C, Jacques M-A, Boureau T (2013) High throughput quantitative phenotyping of plant resistance using chlorophyll fluorescence image analysis. Plant Methods 9: 17

Sapkota S, Mergoum M, Liu Z (2020) The translucens group of Xanthomonas translucens: Complicated and important pathogens causing bacterial leaf streak on cereals. Mol Plant Pathol 21: 291–302

Sapkota S, Zhang Q, Chittem K, Mergoum M, Xu SS, Liu Z (2018) Evaluation of triticale accessions for resistance to wheat bacterial leaf streak caused by Xanthomonas translucens pv. undulosa. Plant Pathol 67: 595–602

Shantharaj D, Römer P, Figueiredo JFL, Minsavage GV, Krönauer C, Stall RE, Moore GA, Fisher LC, Hu Y, Horvath DM, et al (2017) An engineered promoter driving expression of a microbial avirulence gene confers recognition of TAL effectors and reduces growth of diverse Xanthomonas strains in citrus: X. citriTALE-EBE-mediated citrus resistance. Mol Plant Pathol 18: 976–989

Shirley MD, Ma Z, Pedersen BS, Wheelan SJ (2015) Efficient “pythonic” access to FASTA files using pyfaidx. doi: 10.7287/peerj.preprints.970v1

Soneson C, Love MI, Robinson MD (2015) Differential analyses for RNA-seq: transcript-level estimates improve gene-level inferences. F1000Res 4: 1521

Streubel J, Pesce C, Hutin M, Koebnik R, Boch J, Szurek B (2013) Five phylogenetically close rice SWEET genes confer TAL effector-mediated susceptibility to Xanthomonas oryzae pv. oryzae. New Phytol 200: 808–819

Supek F, Bošnjak M, Škunca N, Šmuc T (2011) REVIGO Summarizes and Visualizes Long Lists of Gene Ontology Terms. PLOS ONE 6: e21800

Tameling WIL, Elzinga SDJ, Darmin PS, Vossen JH, Takken FLW, Haring MA, Cornelissen BJC (2002) The tomato R gene products I-2 and MI-1 are functional ATP binding proteins with ATPase activity. Plant Cell 14: 2929–2939

Taniguchi S, Hosokawa-Shinonaga Y, Tamaoki D, Yamada S, Akimitsu K, Gomi K (2014) Jasmonate induction of the monoterpene linalool confers resistance to rice bacterial blight and its biosynthesis is regulated by JAZ protein in rice. Plant Cell Environ 37: 451–461

Teper D, White FF, Wang N (2023) The Dynamic Transcription Activator-Like Effector Family of Xanthomonas. Phytopathology®. doi: 10.1094/PHYTO-10-22-0365-KD

Urzinger S, Würstl L, Avramova V, Urbany C, Scheuermann D, Presterl T, Reuscher S, Mayer M, Brajkovic S, Küster B, et al (2026) Native alleles at lhcb6 shape photosynthetic efficiency and early growth in maize. Sci Rep. doi: 10.1038/s41598-026-42348-8

Wang D, Wei L, Liu T, Ma J, Huang K, Guo H, Huang Y, Zhang L, Zhao J, Tsuda K, et al (2023) Suppression of ETI by PTI priming to balance plant growth and defense through an MPK3/MPK6-WRKYs-PP2Cs module. Mol Plant 16: 903–918

Wang L, Rinaldi FC, Singh P, Doyle EL, Dubrow ZE, Tran TT, Pérez-Quintero AL, Szurek B, Bogdanove AJ (2017) TAL effectors drive transcription bidirectionally in plants. Mol Plant 10: 285–296

Wang Q, Shakoor N, Boyher A, Veley KM, Berry JC, Mockler TC, Bart RS (2021) Escalation in the host-pathogen arms race: A host resistance response corresponds to a heightened bacterial virulence response. PLoS Pathog 17: e1009175

Wen A, Jayawardana M, Fiedler J, Sapkota S, Shi G, Peng Z, Liu S, White FF, Bogdanove AJ, Li X, et al (2018) Genetic mapping of a major gene in triticale conferring resistance to bacterial leaf streak. Theor Appl Genet 131: 649–658

White FF, Potnis N, Jones JB, Koebnik R (2009) The type III effectors of Xanthomonas. Mol Plant Pathol 10: 749–766

Wimalanathan K, Lawrence-Dill CJ (2021) Gene Ontology Meta Annotator for Plants (GOMAP). Plant Methods 17: 54

Xu X, Feng Y, Fang S, Xu J, Wang X, Guo W (2016) Genome-wide characterization of the β-1,3-glucanase gene family in Gossypium by comparative analysis. Sci Rep 6: 29044

Xu Z, Xu X, Li Y, Liu L, Wang Q, Wang Y, Wang Y, Yan J, Cheng G, Zou L, et al (2024) Tal6b/AvrXa27A, a hidden TALE targeting the susceptibility gene OsSWEET11a and the resistance gene Xa27 in rice. Plant Commun 5: 100721

Xu Z-Y, Zou L-F, Ma W-X, Cai L-L, Yang Y-Y, Chen G-Y (2017) Action modes of transcription activator-like effectors (TALEs) of Xanthomonas in plants. J Integr Agric 16: 2736–2745

Zhang H, Tang M, Wan Y, Deng Z, Qin X, Huang J, Wei X, Li R, Liu F (2025) Transcriptome analysis of rice resistant and susceptible near-isogenic lines in response to infection by pv. Front Plant Sci 16: 1610315

