## Supplementary figures and images for "Distinct Temporal Responses to *Xanthomonas translucens* Strains Shape Bacterial Leaf Streak Development in Triticale"

### Supplementary Figure 1

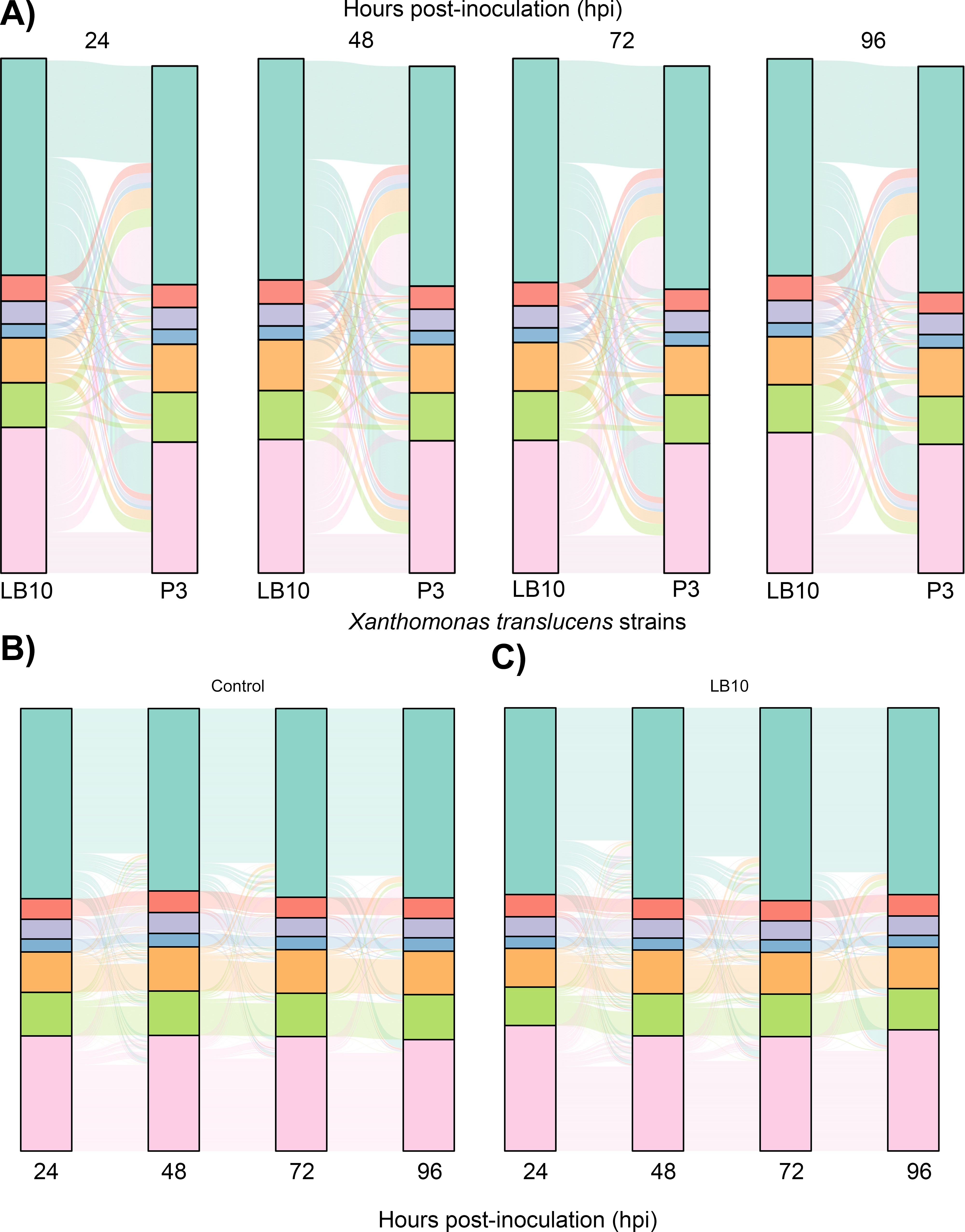

### Supplementary Figure 2

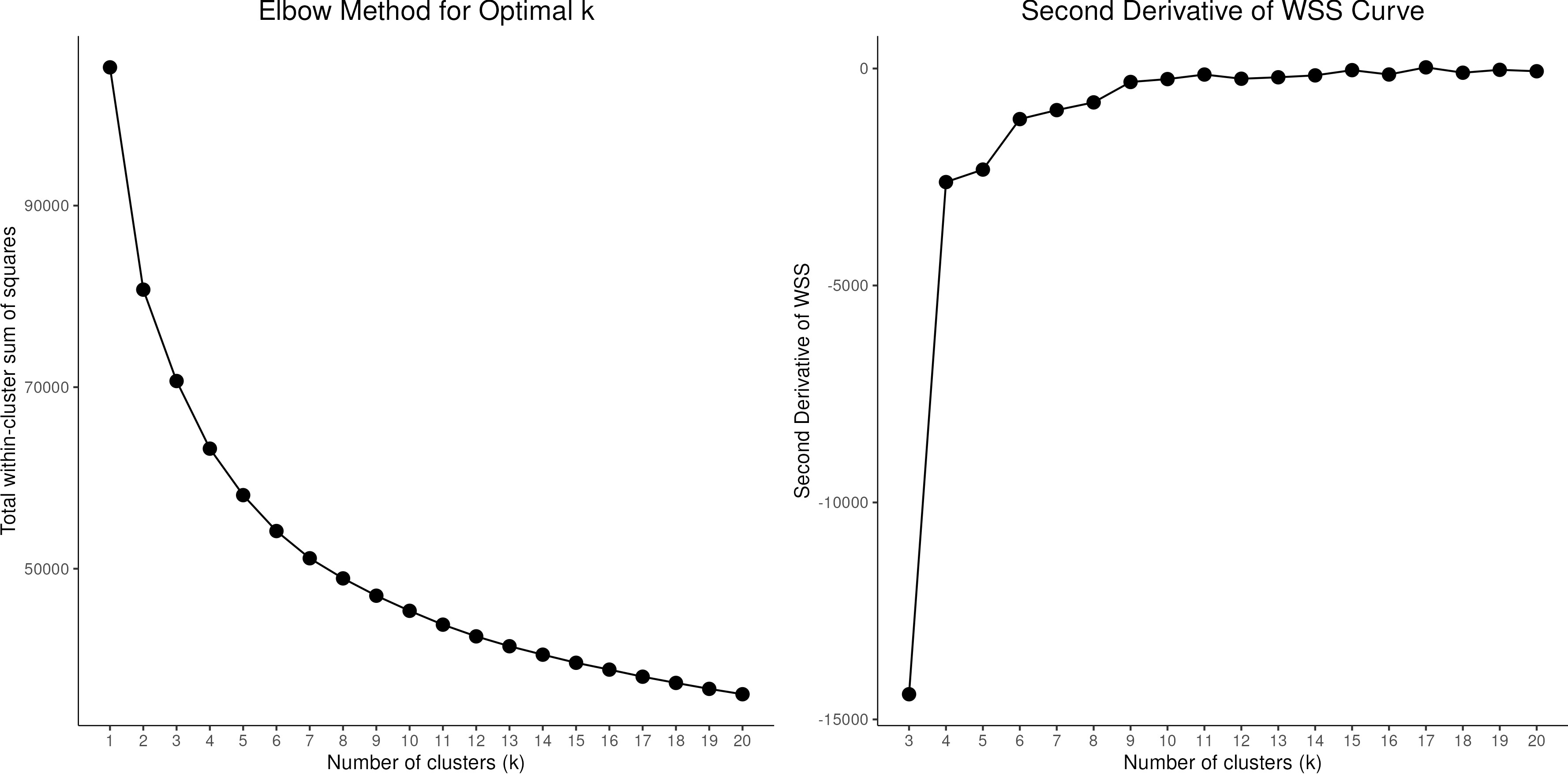

### Supplementary Figure 3

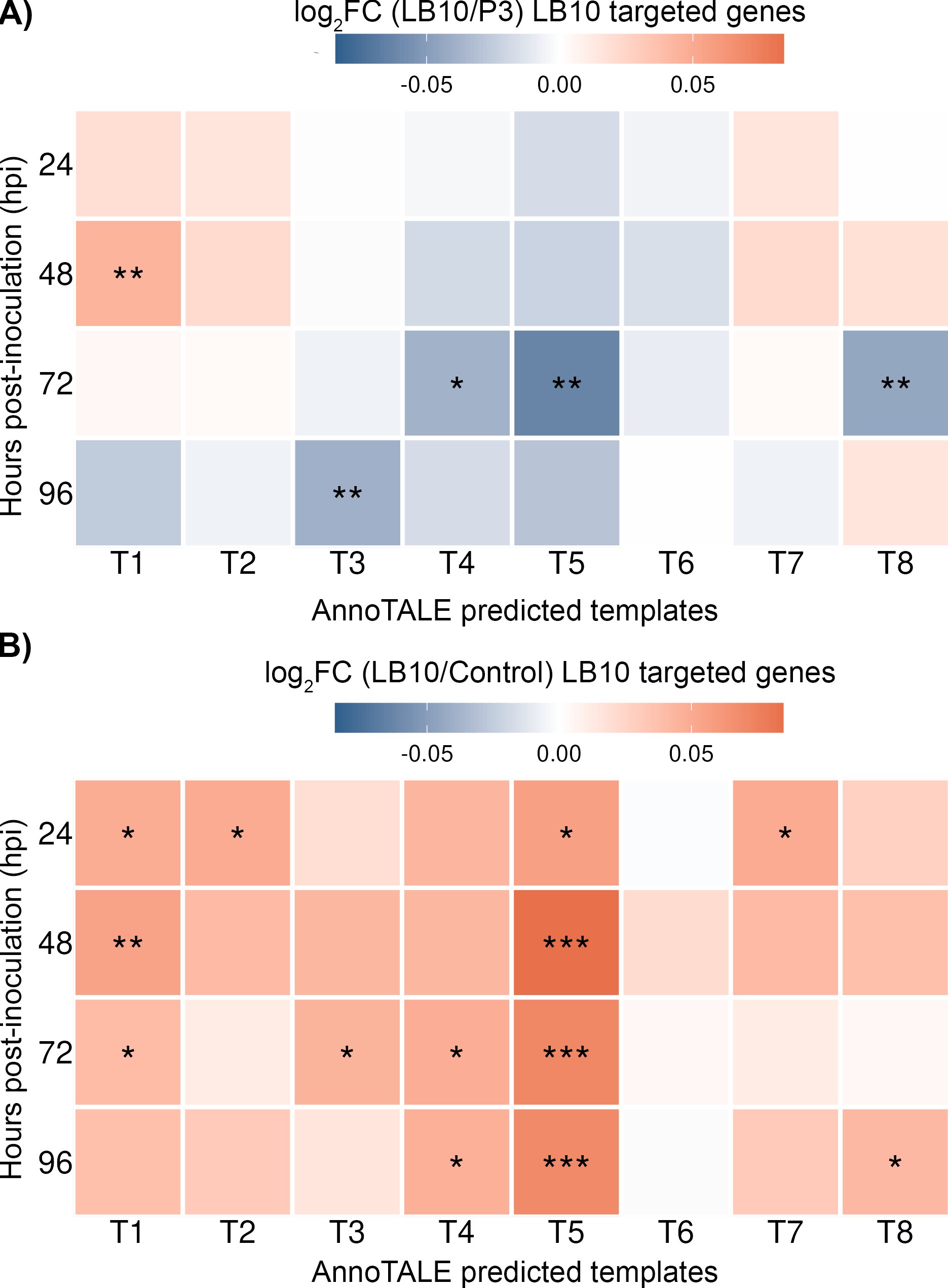

### Supplementary Figure 4

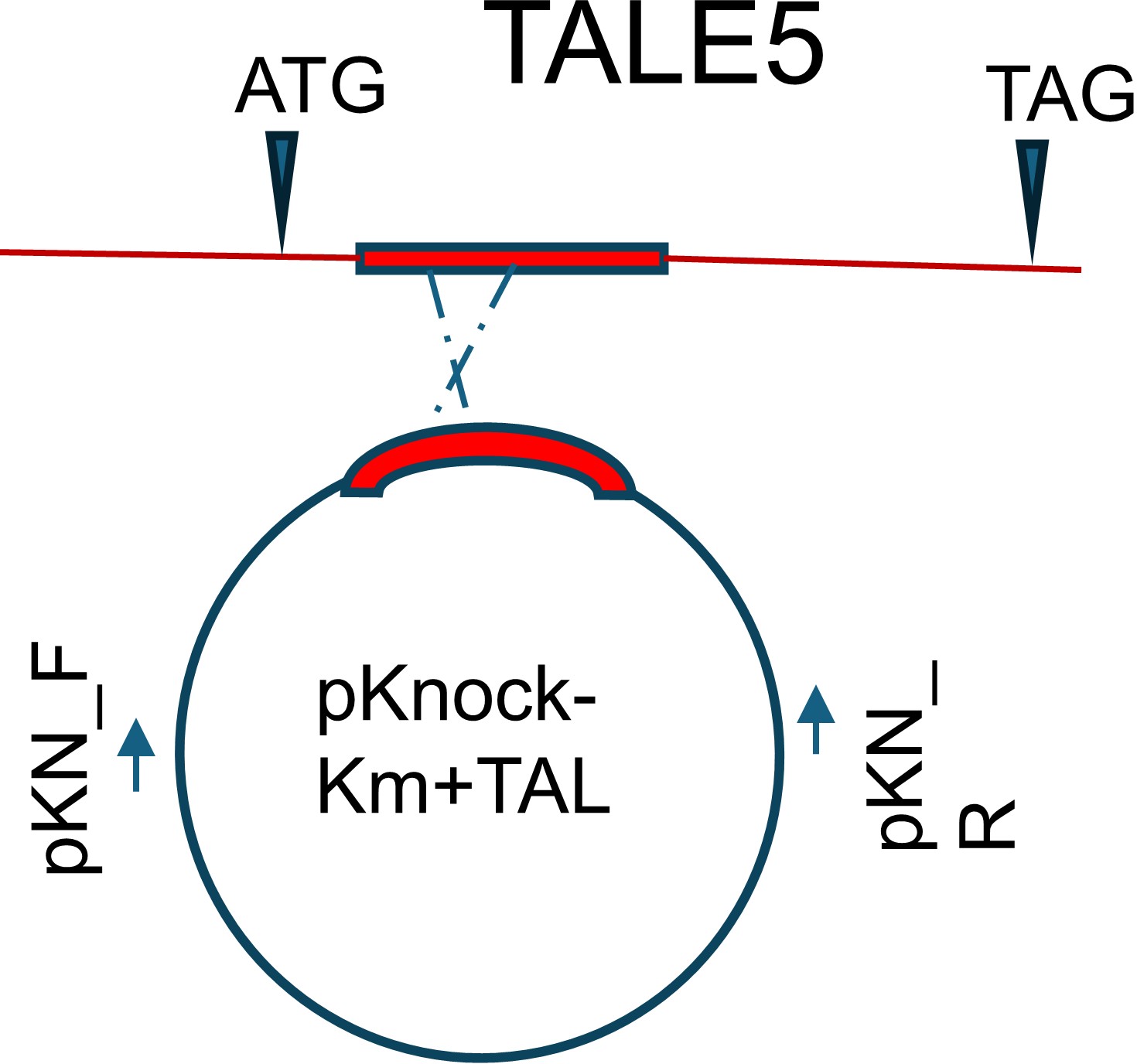

### Supplementary Figure 5

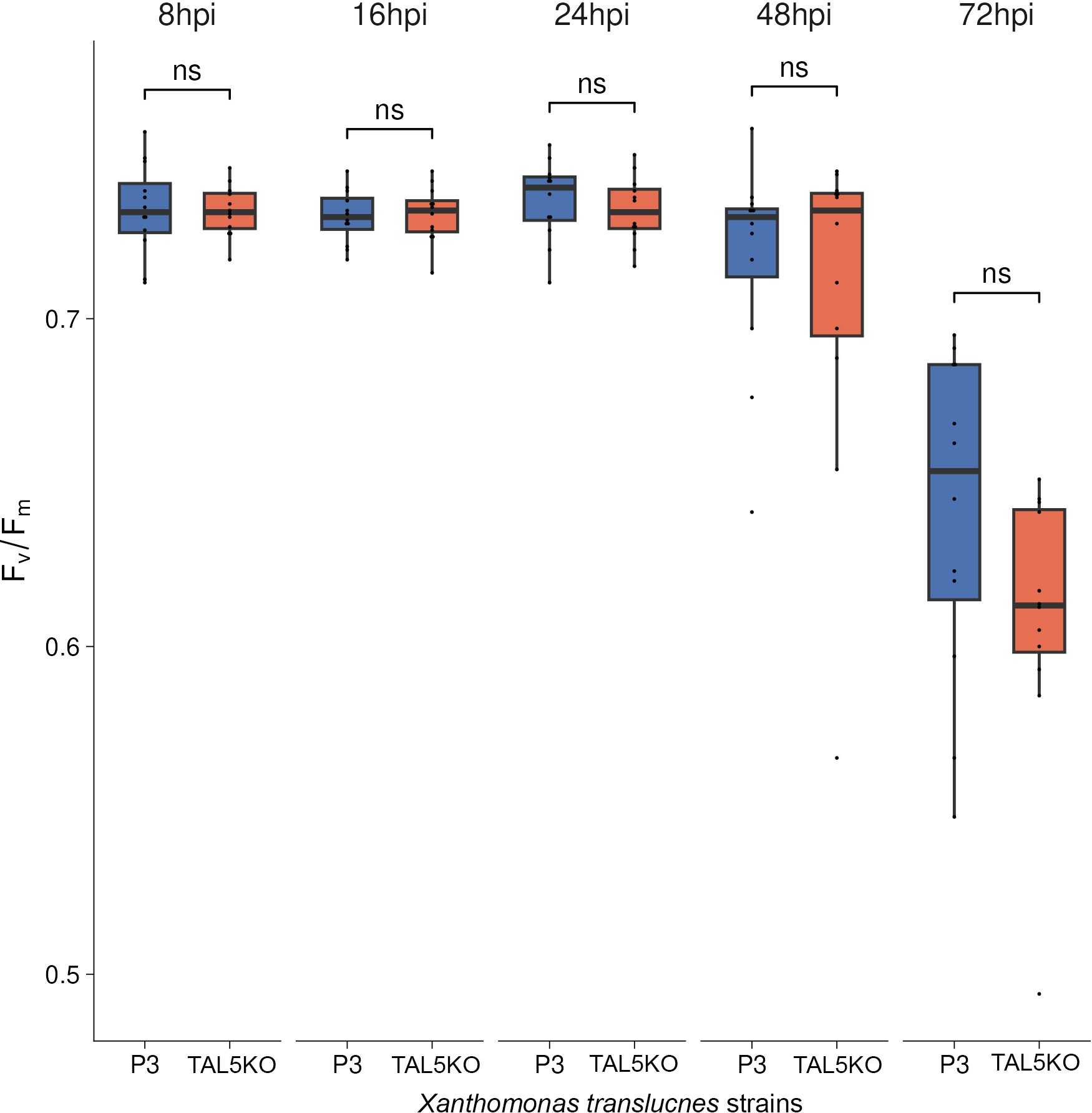
